# A conserved interaction between a C-terminal motif in Norovirus VPg and the HEAT-1 domain of eIF4G is essential for translation initiation

**DOI:** 10.1101/024349

**Authors:** Eoin N. Leen, Frédéric Sorgeloos, Samantha Correia, Yasmin Chaudhry, Fabien Cannac, Chiara pastore, Yingqi Xu, Stephen C. Graham, Stephen J. Matthews, Ian G. Goodfellow, Stephen Curry

## Abstract

Translation initiation is a critical early step in the replication cycle of the positive-sense, single-stranded RNA genome of noroviruses, a major cause of gastroenteritis in humans. Norovirus RNA, which has neither a 5′ m7G cap nor an internal ribosome entry site (IRES), adopts an unusual mechanism to initiate protein synthesis that relies on interactions between the VPg protein covalently attached to the 5′-end of the viral RNA and eukaryotic initiation factors (eIFs) in the host cell.

For murine norovirus (MNV) we previously showed that VPg binds to the middle fragment of eIF4G (4GM; residues 652-1132). Here we have used pull-down assays, fluorescence anisotropy, and isothermal titration calorimetry (ITC) to demonstrate that a stretch of ∼20 amino acids at the C terminus of MNV VPg mediates direct and specific binding to the HEAT-1 domain within the 4GM fragment of eIF4G. Our analysis further reveals that the MNV C-terminus binds to eIF4G HEAT-1 via a motif that is conserved in all known noroviruses. Fine mutagenic mapping suggests that the MNV VPg C terminus may interact with eIF4G in a helical conformation. NMR spectroscopy was used to define the VPg binding site on eIF4G HEAT-1, which was confirmed by mutagenesis and binding assays. We have found that this site is non-overlapping with the binding site for eIF4A on eIF4G HEAT-1 by demonstrating that norovirus VPg can form ternary VPg-eIF4G-eIF4A complexes. The functional significance of the VPg-eIF4G interaction was shown by the ability of fusion proteins containing the C-terminal peptide of MNV VPg to inhibit translation of norovirus RNA but not cap- or IRES-dependent translation. These observations define important structural details of a functional interaction between norovirus VPg and eIF4G and reveal a binding interface that might be exploited as a target for antiviral therapy.

## Introduction

The *Caliciviridae* family of positive-sense, single-strand RNA viruses includes 5 genera that infect a variety of animals: *Norovirus, Lagovirus, Nebovirus, Sapovirus and Vesivirus*. Of these, the noroviruses constitute the greatest threat to human health. They cause around 18% of all cases of acute gastroenteritis worldwide and an estimated 200,000 deaths a year in children under 5 in developing nations [1,2]. The *Norovirus* genus contains 6 genogroups (GI-GVI), three of which infect humans (GI, GII and GIV). The murine noroviruses (MNV) of the GV genogroup only infect rodents and have been studied extensively as a convenient model for human norovirus infections [3-6].

A critical step in calicivirus replication following delivery of the viral RNA to the cytoplasm of infected cells is the initiation of translation. As obligate intracellular parasites, viruses need to gain access to the protein synthesis machinery and, in common with many other viruses, caliciviruses do this by partially circumventing the host mechanisms of translation initiation [7]. The initiation of protein synthesis is a complex and highly regulated process by which the 43 S ribosomal initiation complex is first recruited to an mRNA molecule and then directed to the AUG start codon [8]. In eukaryotes this process involves a multitude of protein initiation factors along with initiator tRNA.

In the normal or canonical mechanism of ribosomal recruitment an important early step is the binding of the heterotrimeric eukaryotic initiation factor 4F (eIF4F) to the m7G cap at the 5′end of the mRNA. eIF4F mediates binding of the mRNA to the 43 S ribosomal initiation complex via its interaction with eIF3 and, in concert with a number of other initiation factors, promotes scanning of the mRNA until the initiation complex locates the AUG start codon [8,9]. At this point ribosomal assembly is completed and protein synthesis commences. eIF4F consists of a large, multi-domain ‘platform’ protein, eIF4G, which has independent sites for recruitment of eIF4E, the cap-binding protein, and eIF4A, an ATP-dependent RNA helicase that melts RNA secondary structure to facilitate ribosomal scanning (Fig 1A). eIF4G also serves as a platform for other proteins such as the polyA-binding protein (PABP), which augments the efficiency of translation initiation, and the MAP kinase-interacting kinases 1 and 2, which are involved in regulation (reviewed in [9]). In humans there are two closely-related paralogs of eIF4G (I and II) which share 46% sequence identity [10]; there are also a number of alternatively spliced isoforms for each paralog but the differentiation of their functions in translation has yet to be fully determined [11]. DAP5 (Death-associated protein 5; also known as p97, NAT-1 and eIF4G2) is another eIF4G paralog [12]. It is missing the N-terminal portions of eIF4GI and eIF4GII which contain the motifs that bind PABP and eIF4E but it otherwise similar in domain structure to the C-terminal two-thirds of these proteins (Fig 1A), sharing 39% and 43% amino acid sequence identity respectively.

**Figure 1.**
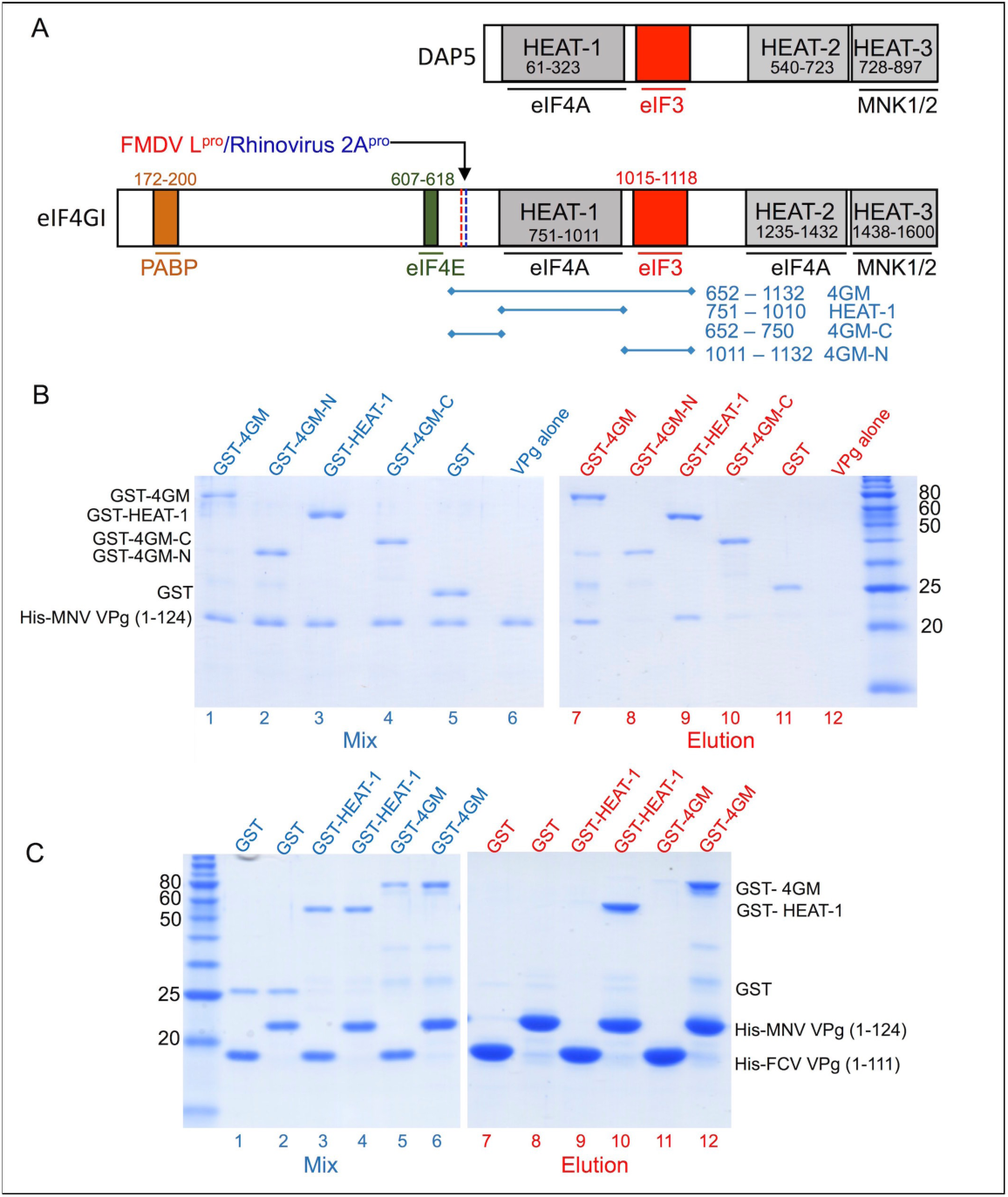
**MNV VPg interacts with eIF4GI via its HEAT-1 domain**. (A) Schematic representation of eIF4GI (NCBI accession AAM69365.1), one of two eIF4G paralogues expressed in humans, and the paralog DAP5 (NCBI accession NP_001036024.3). Positions of domains that interact with other proteins of the translation initiation machinery are indicated, as are the cleavage sites of FMDV L protease and Rhinovirus 2A protease. The principal eIF4G fragments that were sub-cloned for use in this study are also indicated. (B) SDS PAGE -analysis of glutathione affinity pull-down assays that were performed to map the locus of MNV VPg binding. GST-fusions of various eIF4GI fragments (shown in panel A) were used as bait and His-tagged-MNV VPg(1-124) as prey. Left panel: protein mixtures applied to the glutathione-sepharose 4B beads (lanes 1–6). Right panel: proteins eluted with 10 mM glutathione (lanes 7–12). (C) SDS PAGE analysis of cobalt affinity pull-down assays performed to confirm the eIF4G HEAT-1 domain as the locus of MNV VPg binding. His-tagged-MNV VPg 1-124 or His-tagged-FCV VPg 1-111 as bait proteins and GST-fusions of various eIF4GI fragments were used as prey. Left panel: protein mixtures applied to the cobalt resin (Lanes 1–6). Right panel: proteins eluted with 250 mM imidazole (lanes 7–12).

The adoption of non-canonical mechanisms of ribosomal recruitment is used by some RNA viruses to gain an advantage over host cell mRNA in competing for ribosome recruitment. A common strategy is the use of an internal ribosome entry site (IRES), a large RNA structure within the 5′ untranslated region (5′UTR) of the viral genome, which can mediate ribosome binding independently of a 5′cap and which typically has a reduced dependency on eIFs for translation initiation [13]. Viruses in possession of an IRES can therefore gain a competitive advantage by disabling redundant eIFs that are critical for cap-dependent translation initiation [7,14-17]. Some of the best-studied examples of this strategy are to be found among the picornaviruses (*e.g.* poliovirus (PV), human rhinovirus (HRV), and foot-and-mouth disease virus (FMDV)) [18-20]. Their RNA genomes lack a 5′cap and encode proteases (PV 2A^pro^, HRV 2A^pro^, FMDV L^pro^) that can specifically cleave eIF4G to separate the binding sites for eIF4E and eIF4A [21]. This disables cap-dependent translation initiation, leading to shut-off of host cell protein synthesis, but does not impair IRES-dependent translation initiation from the viral genome, which only requires the large C-terminal fragment of eIF4G retaining the binding site for eIF4A [9]. Consistent with its inability to bind the cap-binding protein eIF4E, DAP5 has been reported to preferentially support IRES-dependent translation initiation [22].

Although picornaviruses and caliciviruses share many similarities in their genome structures and replication cycles, caliciviruses have acquired completely different mechanism for translation initiation. They have no IRES – indeed their 5′UTRs are very small, in some cases as short as 4 nucleotides (nt) [5,23] – but instead rely on a small protein called VPg (Virus Protein, genome-linked) of ∼110-140 amino acids that is covalently linked to the 5′end of the viral genome for translation initiation. Picornaviruses are also known to have a VPg attached to the 5′ends of their RNA genomes but the picornaviral protein, which is typically only around 22 amino acids and structurally unrelated to its caliciviral equivalent, is rapidly removed following cell entry [24] and plays no part in translation [25,26]. In contrast, proteolytic digestion of caliciviral VPg *in vitro* has long been known to render its RNA non-infectious and to prevent translation *in vitro* [27-30]. The observation that infectivity of caliciviral RNA can be restored by covalent attachment of a 5′ m7G cap suggests that VPg serves as a proteinaceous cap analogue [30,31]. However, the particulars of the mechanism by which VPg performs this role have yet to be fully elucidated.

Picornavirus VPg peptides appear to be largely unstructured in aqueous buffer [32], but recent work of the solution structures of caliciviral VPg proteins from three different genera has revealed that they all contain a helical core flanked by unstructured N and C termini. The feline calicivirus (FCV, a vesivirus) and porcine sapovirus (PSaV) VPg cores consist of very similar 3 helix bundles, while the core of MNV VPg is somewhat truncated and contains just the first two helices found in the other two proteins [33,34].

These structural similarities belie the fact that, despite being critical for translation initiation, VPg proteins from different caliciviruses appear to perform this function in different ways. One of the earliest studies found that the VPg of noroviruses interact with eIF3 but the mechanistic implications of this finding remain unclear [35,36]. Direct interactions with eIF4E have been reported for FCV VPg and MNV VPg [37] but, while this interaction appears to be crucial for FCV, its significance for MNV is more questionable. For example, disruption of the eIF4E-eIF4G interface by the translational regulator 4E-BP1 inhibits FCV translation, but has no effect on MNV translation [27]. Moreover, FCV translation is abrogated if FMDV L^pro^ is allowed to digest eIF4G, thereby separating its eIF4E and eIF4A binding domains, but MNV translation is unaffected by this proteolytic cleavage. These findings reinforce the idea that FCV VPg probably operates as a functional analogue of the m^7^G cap, by interacting directly with eIF4E, whereas MNV relies on a distinct mode of interaction with eIF4F (see below). More recent work suggests that PSaV VPg may direct translation initiation by a mechanism similar to FCV VPg since it also binds directly to eIF4E [38]; moreover, PSaV translation *in vitro* also exhibits the same sensitivity to inhibition by 4E-BP1 and L^pro^ digestion of eIF4G.

The translation initiation directed by MNV VPg may not be completely independent of eIF4E in view of a recent study showing that modulation of the MAPK pathway to stimulate phosphorylation of eIF4E during MNV1 infection helps to promote virus replication [17]. Nevertheless it is clear that the interaction of MNV VPg with eIF4E is not its primary mode of engagement with the translational machinery. Recent work revealed that MNV VPg is instead capable of forming a specific interaction with eIF4G [39]. The interaction was identified initially using tandem affinity purification of double-tagged MNV VPg to identify specific binding partners in eukaryotic cell lysates [39]. The direct nature of the interaction was confirmed using purified recombinant proteins. The site of VPg interaction was mapped to the central 4GM fragment of eIF4G (residues 652-1132) and alanine scanning mutagenesis of VPg suggested that residues in the C terminus made important contributions to binding affinity [39].

The work described here extends these initial observations by providing a detailed characterisation of the interaction of MNV VPg with eIF4G. We describe a biochemical and biophysical dissection of the interaction using a combination of qualitative and quantitative binding assays, and NMR analyses to map precisely the particular regions of MNV VPg and eIF4G that mediate binding. We have discovered a C-terminal sequence motif in MNV VPg that is conserved in all noroviruses and is capable of recapitulating the interactions made by MNV VPg with the translation initiation machinery in living cells. We further find that this motif binds to the HEAT-1 domains of eIF4GI and eIF4GII with micromolar affinity. Our evidence suggests that this VPg peptide motif may adopt a helical conformation when bound to eIF4G HEAT-1 and can mediate the formation of a ternary VPg-eIF4G-eIF4A complex with 1:1:1 stoichiometry in the presence of eIF4A. The functional significance of this interaction is evident from the observation that peptides corresponding to the eIF4G-binding motif of MNV VPg mediate an interaction with translation initiation complexes in cells that is indistinguishable from that made by the intact protein, and inhibit norovirus translation *in vitro*. Further our findings provide proof of principle that the VPg-eIF4G interaction might provide a useful target for antiviral drug development. The observation that the VPg-NS6^pro^ precursor is unable to bind eIF4G shows that a free VPg C terminus is required for the interaction and suggests that proteolytic processing of the viral polyprotein precursor may be linked to the differential regulation of translation and RNA replication phases in the virus life cycle. Taken together our results provide significant new insights into the molecular mechanism of norovirus translational initiation mechanisms of noroviruses and identify an interaction interface that could be targeted for antiviral drug development.

## RESULTS

### MNV VPg binds to the eIF4G 4GM region via the eIF4G HEAT-1 domain

We previously showed that the binding site on eIF4GI for MNV VPg resides within the central 4GM fragment (residues 652 -1132; Fig 1A) [39]. This 4GM fragment contains a central HEAT-1 domain (residues 751-1010), a crescent-shaped module comprised of 10 α-helices [40], flanked by N- and C-terminal regions that are not known to be structured. To determine whether MNV VPg bound to the HEAT-1 domain we prepared GST-fusion proteins corresponding to sub-fragments of 4GM – the N- and C-terminal flanking regions (GST-4GM-N and GST-4GM-C respectively), and GST-eIF4G-HEAT-1 (Fig 1A) – and used them as bait proteins in glutathione pull-down assays to test their ability to interact with the viral protein (Fig 1B). These assays showed that GST-4GM and GST-eIF4G-HEAT-1 bound His-tagged MNV VPg with similar efficiency (lanes 7, 9) but that neither GST-4GM-N nor GST-4GM-C (or GST alone) had any detectable affinity for VPg (lanes 8, 10, 11).

In a second set of pull-down assays, performed on cobalt affinity resin using His-tagged MNV VPg as the bait, the same pattern of binding was observed (see (Fig 1C)): GST-4GM and GST-eIF4G-HEAT-1 bound His-tagged MNV VPg equally well (lanes 10, 12) but there was no observed interaction with GST alone (lane 8). This experiment also showed that FCV VPg, which is only 27% identical in amino acid sequence with MNV VPg, does not interact with either GST-4GM or GST-HEAT-1 (lanes 9, 11).

Together these results suggest that MNV VPg interacts with the HEAT-1 domain of eIF4GI.

### A conserved sequence at the C terminus of MNV VPg is critical for HEAT-1 domain binding

Previous work identified three mutations close to the C terminus of the 124-residue MNV VPg, V115A, D116A and F123A, which all reduced its interaction with eIF4F [39]. This observation is likely due to the effect of the mutations on binding to eIF4G since the F123A mutation was also shown to substantially reduce binding of VPg to GST-4GM. The possible functional importance of the C terminus of MNV VPg was further underscored by alignment of VPg amino acid sequences from representative members of each of the norovirus genogroups, which revealed that residues 108-124 are highly conserved (Supplementary Fig S1). Since our NMR studies of MNV VPg have shown that the C-terminal half of MNV VPg (residues 63-124) is flexible and exhibits no interaction with the structured core of the protein [34], the sequence conservation in this region probably reflects the functional constraint imposed by the interaction with eIF4G. We therefore hypothesized that VPg is likely to bind to eIF4G HEAT-1 via a contiguous sequence close to the C terminus.

As a first test of this hypothesis, we compared the ability of MNV VPg proteins with and without the unstructured C terminus to bind to eIF4G HEAT-1. Using GST-eIF4GI HEAT-1 containing a C-terminal His-tag as the bait protein in a cobalt affinity pull-down assay, we found that it could interact with untagged MNV VPg(1124) but not with MNV VPg(185), a truncated version of the protein that retains the helical core but lacks the unstructured C terminus (Fig 2A). This confirms that binding to eIF4G requires the C terminus of VPg.

**Figure 2.**
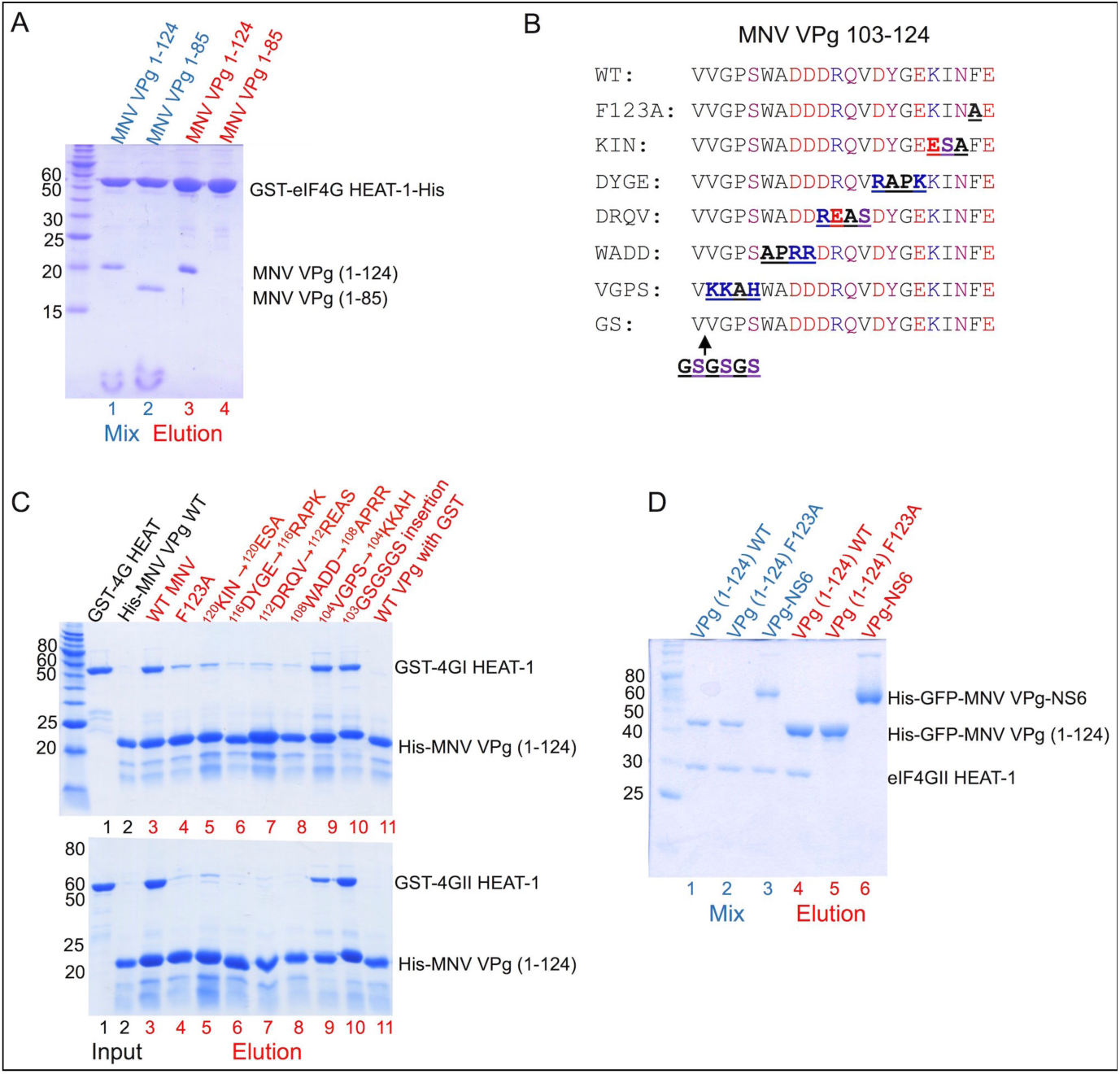
**MNV VPg interacts with eIF4GI and eIF4GII HEAT-1 domains via the C-terminal residues 104-124.** (A) SDS PAGE analysis of cobalt affinity pull-down assay using a GST-eIF4GI HEAT-1 construct with a C-terminal His-tag as bait and either untagged MNV VPg(1-124) or MNV VPg(1-85) as prey. Protein mixtures are shown in lanes 1 and 2 (blue labels); bound proteins eluted with 250 mM imidazole are in lanes 3and 4 (red labels). (B) Sequence alignment showing the location of amino acid substitutions introduced into the C terminus of His-tagged MNV VPg. (C) SDS PAGE analysis of cobalt affinity assays using His-tagged MNV mutants (panel B) as bait and either GST-eIF4GI-HEAT-1 (top panel) or GST-eIF4GII-HEAT-1 as prey. Lanes 1-2: input proteins; lanes 3-11: eluted proteins. (D) SDS PAGE analysis of cobalt affinity pull-down assay using either His-tagged GFP-VPg(1-124) wild-type, GFP-VPg(1-124) F123A or a His-tagged GFP-VPg-NS6 fusion (containing the inactivating C139A mutation of the protease active site Cys (NS6 numbering)) as bait, and untagged eIF4GII HEAT-1 as prey. Lanes 1-3: protein mixes; lanes 4-6: protein eluted with 250 mM imidazole.

To map the extent of the MNV VPg C terminus needed to bind eIF4G we introduced blocks of 3-4 amino acid substitutions within and beyond the conserved sequence motif. In addition to the F123A mutation already known to reduce binding to eIF4G [39], we made the following mutations in His-tagged MNV VPg: ^120^KIN→^120^ESA, ^116^DYGE→^116^RAPK, ^112^DRQV→^112^REAS, ^108^WADD→^108^APRR, ^104^VGPS→^104^KKAH, and an insertion of GSGSGS after residue 103 (Fig 2B). Wild-type and mutant MNV VPg proteins were used as bait proteins in cobalt affinity pull-down assays to determine their ability to bind GST-eIF4GI and GST-eIF4GII HEAT-1 proteins, which are 83 % identical in amino acid sequence within this domain (Fig 2C). All mutations made in the conserved region (residues 108-123) had drastically reduced binding capacity for eIF4GI HEAT-1. However, neither the ^104^VGPS→^104^KKAH mutations nor the GSGSGS insertion at position 103, which introduced substitutions at the N-terminal end of the conserved motif, had any observable effect on MNV VPg binding to eIF4GI HEAT-1 (Fig 2C, upper panel). A very similar pattern of effects was observed when the MNV VPg mutants were tested for binding to GST-eIF4GII HEAT-1, with the exception that the ^104^VGPS→^104^KKAH mutations were found to moderately reduce binding of the viral protein (Fig 2C, lower panel). These data are consistent with the hypothesis that the conserved motif within residues 108-124 of MNV VPg plays a critical role in eIF4G binding, and that the N-terminal boundary of the binding domain for eIF4GI and eIF4GII lies between residues 104 and 108.

Finally, given that the eIF4G HEAT-1 binding site on MNV VPg is located very close to the C terminus of the protein, which is generated by proteolytic processing at the VPg-NS6 junction within the virus polyprotein precursor, we asked whether the interaction with eIF4G HEAT-1 requires a free C terminus. To address this question we performed cobalt affinity pull down assays to compare the ability of eIF4GII HEAT-1 to bind wild-type his-GFP tagged MNV VPg and a his-GFP tagged MNV VPg-NS6 fusion protein which contained a C139A mutation in the NS6 protease to prevent auto-processing of the VPg-NS6 junction; (the GFP tags are not relevant to this assay – they were added in anticipation of using these reagents for other experiments that were not performed). We found that wild-type his-GFP-VPg was effective at capturing eIF4GII HEAT-1 in these assays but his-GFP-VPg-NS6 had no observable binding activity (Fig 2D). The fusion protein exhibited the same lack of binding activity as the his-GFP-VPg (F123A) mutant, which was used as a negative control (Fig 2D). This result suggests that processing of the VPg-NS6 junction is required for the MNV VPg-eIF4G interaction.

### The C terminus of MNV VPg binds eIF4G HEAT-1 domains with micromolar affinity

The results described above indicate that a contiguous region at the C terminus of VPg mediates binding to the eIF4G HEAT-1 domain. To determine the affinity of the interaction, we used peptides corresponding to the MNV VPg C terminus in fluorescence anisotropy experiments. Peptides corresponding to MNV VPg 104-124 and 108-124 and incorporating an N-terminal FITC group were chemically synthesized and tested for binding to untagged versions of the HEAT-1 domains of eIF4GI (748-993) and eIF4GII (751-1003) (see Materials and Methods). In addition, we tested the binding of these peptide to the HEAT-1 domain of DAP5 (61-323), which shares approximately 40% amino acid sequence identity with the HEAT-1 domains of eIF4GI and eIF4GII and is very similar in structure [41].

In a first set of experiments, we compared binding of the two MNV VPg peptides to eIF4GII HEAT-1. These showed that MNV VPg 104-124 binds with a K_D_ of 2.8 µM, while the shorter MNV VPg 108-124 peptide binds with about four-fold weaker affinity (K_D_ = 10.6 µM) (Fig 3A). Very similar affinities for these two peptides were observed in experiments performed with the fusion protein MBP-eIF4GII HEAT-1 (Supplementary Fig S2). These results are consistent with the pull-down experiments, which found that mutations on the C-terminal side of position 104 in VPg (^104^VGPS→^104^KKAH) reduced binding to eIF4GII, whereas the ^103^GSGSGS insertion immediately before position 104 did not (Fig 2C).

**Figure 3.**
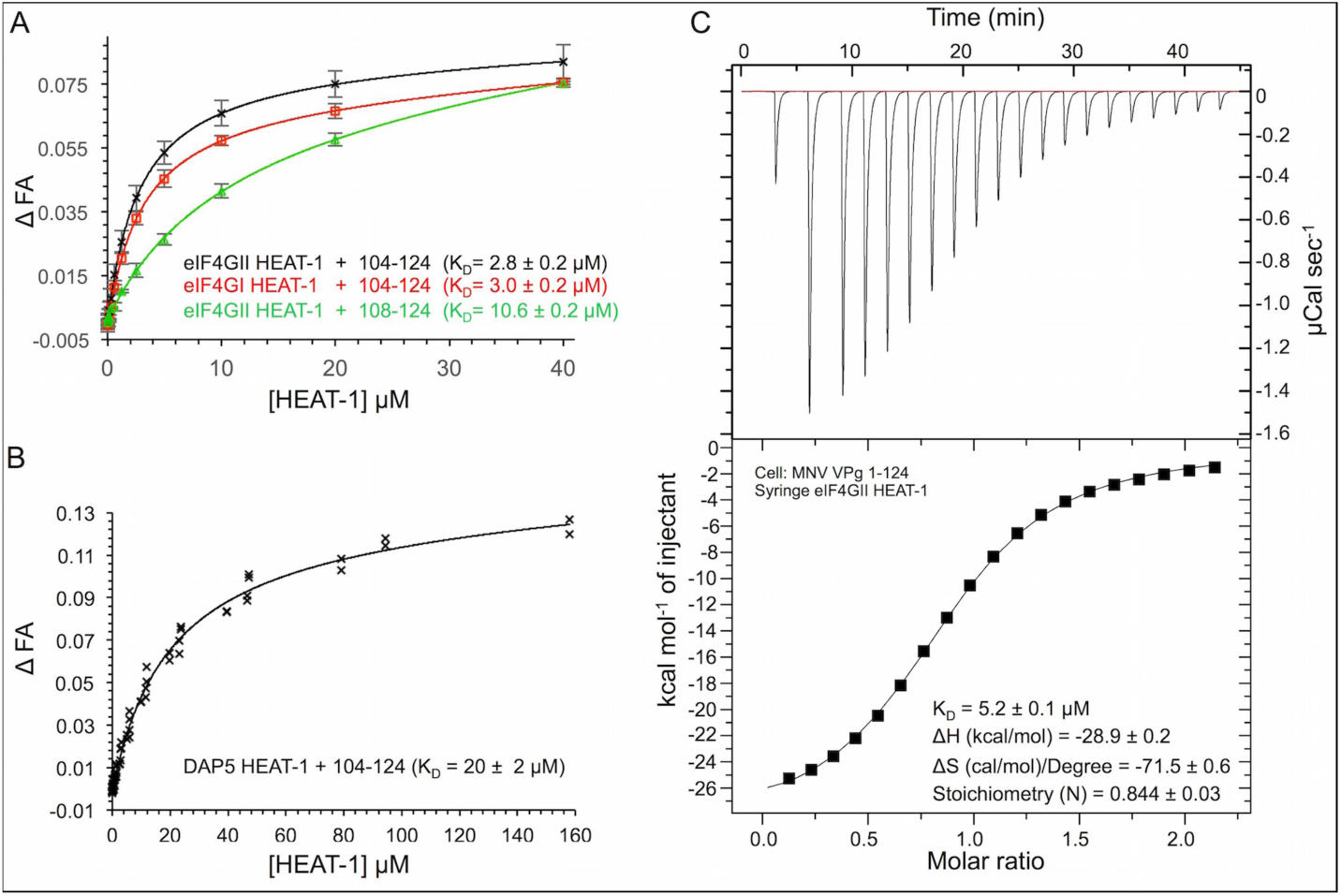
**Figure 3: MNV VPg 104-124 interacts with the HEAT-1 domains of eIF4GI, eIF4GII and DAP5 with low micromolar affinity.** (A-B) FITC-labelled peptides – MNV VPg(104-124) and MNV VPg(108-124) – were used in fluorescence anisotropy binding assays with unlabelled HEAT-1 domains of eIF4GI (748-993), eIF4GII (745-1003) and DAP5 (61-323) in order to measure the affinity of the interaction. The normalised change in fluorescence anisotropy (relative to a no-protein control), ΔFA, is plotted against protein concentration. Error bars in ΔFP indicate the standard deviation of 5 (eIF4GI) or 10 (eIF4GII) independent measurements. The solid lines indicate the fit to a single-site binding model. (A) Comparison of the binding of MNV VPg(104-124) and MNV VPg(108-124) to eIF4GI and eIF4GII HEAT-1 domains. (B) Binding of MNV VPg(104-124) to the HEAT-1 domain of DAP5. The fit is calculated with the fluorescence anisotropy data from three independent experiments (all included in the graph). (C) ITC experiments in which unlabelled MNV VPg 1-124 was titrated into eIF4GII (745-1003). Top panel: raw data obtained for a representative experiment from 20 injections (firstly with a volume of 0.5 µL followed by 19 injections of 2 µL of eIF4GII HEAT-1). Bottom panel: the integrated data with a best-fit curve for the representative experiment generated for a single-site binding model using the Origin software package.

All subsequent binding experiments were performed with the tighter-binding MNV VPg 104-124 peptide. It was observed to bind with essentially identical affinity to the HEAT-1 domains of eIF4GII (K_D_ = 2.8 µM) and eIF4GI (K_D_ = 3.0 µM) (Fig 3A). However MNV VPg(104-124) bound about six-fold less well to the DAP5 HEAT1 domain K_D_ = 19.9 µM) (Fig 3B). This difference in affinity suggests that VPg is more likely to recruit eIF4GI and eIF4GII in infected cells than DAP5 and may explain why DAP5 was not observed as a ligand for MNV VPg in the tandem-affinity purifications that were used originally to identify eIF4G as a direct ligand in cell lysates [39].

To determine the binding affinity of full-length MNV VPg for eIF4GII HEAT-1 interaction we used isothermal titration calorimetry (ITC). In these experiments, small volumes of untagged eIF4GII HEAT-1 were injected into a 200 µL sample of untagged MNV VPg(1-124); the heat changes arising from the interaction of the two proteins were measured and fitted to a single-site binding model (Fig 3C). The K_D_ determined from three independent ITC experiments was 5.2 µM. This value is comparable to the K_D_ obtained from fluorescent anisotropy measurements performed with the MNV VPg(104-124) peptide (2.8 µM) and strongly suggests that MNV VPg residues outside the C-terminal region do not contribute significantly to eIF4G binding.

### The direct interaction between VPg and eIF4GI HEAT-1 is conserved in noroviruses

Having established that the C terminus of MNV VPg could mediate binding of the protein to eIF4G HEAT-1 domains with micromolar affinity, we sought to test how well this function is conserved in related viruses. This seemed likely since 10 out of the 21 residues in the region defined by MNV VPg 104-124 are strictly conserved in all norovirus genogroups (Supplementary Fig S1). Intriguingly, there also appears to be significant conservation of the eIF4G-binding motif in the C terminus of human astrovirus 4 – 11 out of 23 residues are identica; – while there is very little indication of the occurrence of the motif in the C terminus of VPg from FCV, a member of the *Vesivirus* genera of the *Caliciviridae* family (Fig 4A). To investigate the eIF4G-binding properties of these various VPg proteins we fused C-terminal sequences corresponding to the sequences that aligned with residues 102-124 from MNV VPg to GST (Fig 4A). The resulting GST-VPg fusion proteins included C-terminal VPg sequences from norovirus genogroups GI, GII, GIII and GV (MNV), from FCV F9 and from human astrovirus 4 (Fig 4A) and were used as prey in cobalt affinity pull-down assays to determine whether they bound to His-eIF4GI HEAT-1, which was used as bait. All of the norovirus VPg GST fusions were found to interact with eIF4GI HEAT-1, similarly effectively (lanes 4-7), while neither FCV nor human Astrovirus 4 GST-VPg fusions (lanes 8, 9) were observed to bind (Fig 4B). This strongly suggests that direct binding to eIF4G is a conserved function of the C terminus of norovirus VPg proteins. The interaction appears to be specific since partial conservation of the eIF4G-binding motif in human astrovirus 4 VPg was not sufficient to permit binding. The negative result for FCV VPg is consistent with previous work which has shown that it interacts directly with eIF4E and that, in contrast to MNV, FCV translation initiation is inhibited in the presence of the eIF4E regulatory protein, 4E-BP1 [27,37].

**Figure 4.**
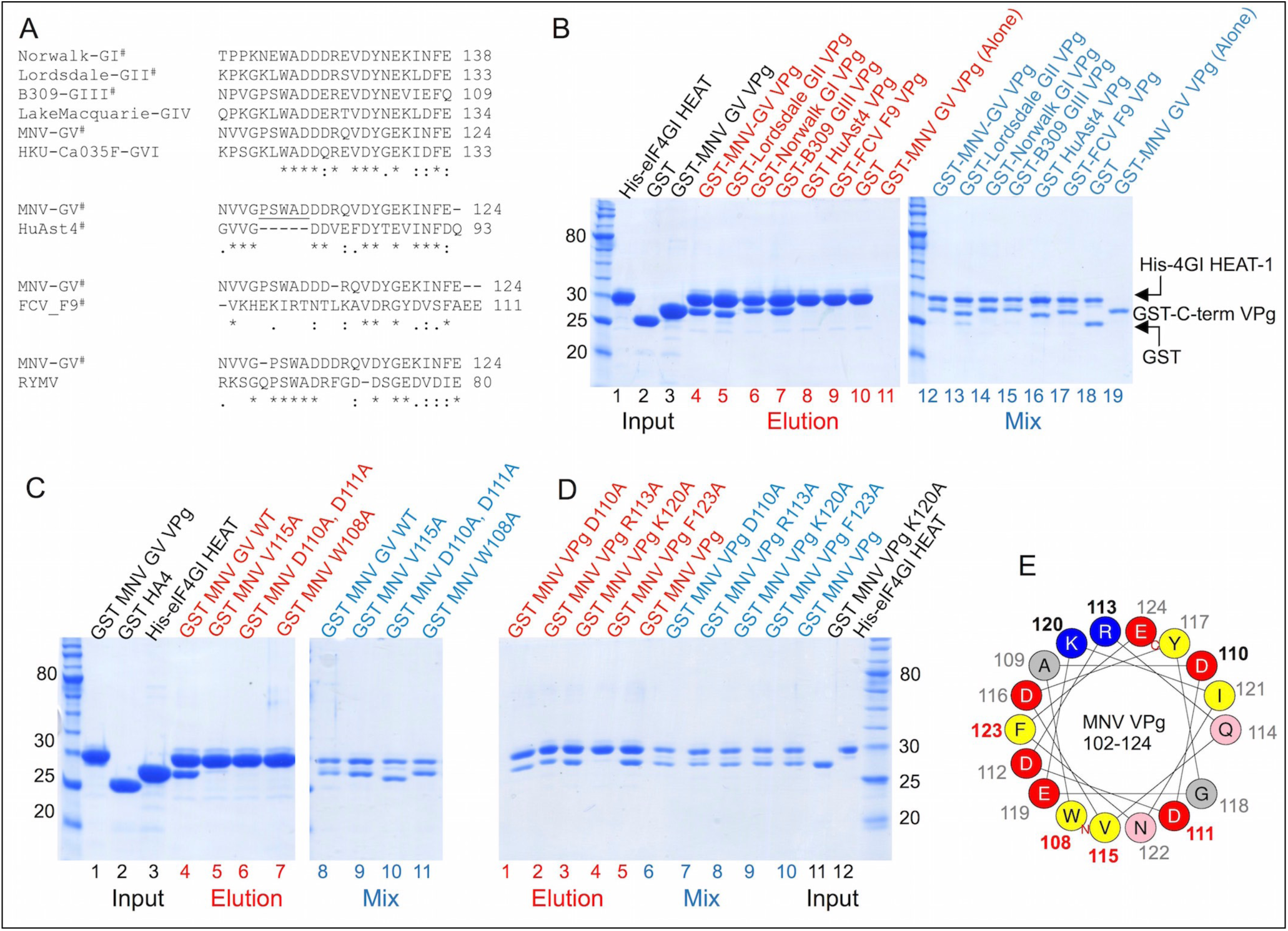
**Figure 4: The C-termini of Noroviral VPg proteins contain a conserved sequence motif that binds to eIF4G HEAT-1.** (A) Top: Amino acid sequence alignments of representative sequences of the C-termini of all 6 genogroups of Norovirus (GI-GIV). Second from top: Alignment of the C-termini of Human Astrovirus 4 (HuAST4) VPg with MNV (GV) VPg. Third from top: Alignment of the C-termini of FCV VPg with MNV (GV) VPg. Bottom: Alignment of the C terminus of MNV (GV) VPg with Rice Yellow Mottle Virus (RYMV) VPg. The representative strains used in the alignments are GI - Hu/GI/Norwalk/1968/US, (NCBI accession AAC64602), GII - Lordsdale virus Hu/GII/Lordsdale/1993/UK (NCBI accession P54634), GIII - Bo/GIII/B309/2003/BEL (NCBI accession ACJ04905.1), GIV - Hu/GIV.1/LakeMacquarie/NSW268O (NCBI accession AFJ21375), GV - Mu/NoV/GV/MNV1/2002/USA (NCBI accession ABU55564.1), GVI - dog/GVI.1/HKU_Ca035F/2007/HKG (NCBI accession FJ692501), FCV - F9 strain (NCBI accession P27409.1), HuAst4 - Human astrovirus 4 (NCBI accession Q3ZN07), Rice yellow mottle virus isolate CI4 (NCBI accession NC_001575). Sequence alignments were performed by ClustalW [6] and BioEdit (http://www.mbio.ncsu.edu/BioEdit/bioedit.html). ^#^Denotes sequences that were used in pull-down assays (see panel B). (B) The indicated VPg sequences in panel A were fused to the C terminus of GST for use as prey in cobalt affinity pull-down assays in which His-tagged eIF4GI HEAT-1 was used as bait, and analysed by SDS PAGE. Lanes 1-3: input samples of some the proteins used in pull-down experiments (black labels); lanes 4-11: eluted proteins (red labels; bands at ∼27 kDa indicate GST-VPg C-terminal constructs that bound to His-eIF4G HEAT-1); lanes 12-19 (blue labels) – protein mixes used in the pull-down experiments. (C, D) Mutational analysis of the C-terminal sequences of MNV VPg were performed using GST-MNV VPg C-terminal fusions in the same way as the experiment presented in panel B. Labelling is colour-coded as in panel B. (E) Helical wheel representation of MNV VPg 102-124 generated using http://heliquest.ipmc.cnrs.fr/ [42]. Residues mutated for the pull-down assays are indicated by bold-face labels and colour-coded by the effect of the substitution on binding to eIF4G HEAT-1: red – no binding; black –binding at or near wild-type levels.

### MNV C terminus may adopt a helical conformation when bound to eIF4G HEAT1

To begin to probe the structural basis of the interaction between MNV VPg and eIF4G HEAT-1, we used GST-fusions with MNV VPg 102-124 to test the contribution to binding of specific amino acids in the C terminus of VPg. In an initial series of experiments we mutated conserved residues within the motif that is responsible for binding to eIF4G; these included W108A, V115A and the double-mutation D110A-D111A. The effects of the mutations were tested using His-eIF4GI HEAT-1 as the bait protein in a pull-down assay. In each case the mutation was found to abrogate binding (Fig 4C; lanes 5-7).

The results of these mutagenesis experiments, along with the previous observation that the F123A point mutation also severely reduces binding (Fig 2C), were intriguing since they suggested that the binding activity is distributed throughout the conserved motif within residues 108-124 of MNV VPg. Since this motif is 17 amino acids long and is found in an unstructured C terminus in the free protein [34], the fact that single point mutations within it (W108A, V115A, F123A) all severely reduced binding suggested that the interaction with eIF4G is unlikely to involve the C terminus of VPg lying in an extended conformation across the surface of the HEAT-1 domain. For such a mode of binding, individual point mutations would be unlikely to completely abrogate binding, particularly for mutations at the extremities of the motif. However, the strong effects of point mutations might be rationalised if the MNV VPg C terminus adopts a rigid structure upon interaction with eIF4G. Consistent with this idea, a helical wheel representation of the MNV VPg C terminus suggests that the mutations tested in the first series of experiments described above (and F123A), would all be on the same side of an α-helix (Fig 4E). This model suggests that there would be an interacting flank (containing W108, V115 and F123) and a non-interacting flank. It also predicts that the reduction in binding observed for the D110A-D111A double mutation is likely to be attributable to the effects of the D111A substitution since it lies on the interacting flank of the helix.

To test the possibility that the C terminus of VPg may adopt a helical conformation upon interacting with eIF4G HEAT-1, we prepared a second set of mutations in the context of our GST-MNV VPg (102-124) fusion protein for use in pull-down assays: D110A, R113A and K120A. Although these mutations are distributed throughout the conserved eIF4G-binding motif in MNV VPg, they were all predicted to be on the non-interacting flank of the putative helix. As can be seen in Fig 4D, GST-MNV VPg (102-124) fusions containing D110A, R113A, and K120A mutations are all capable of binding eIF4GI HEAT-1 (lanes1-3). Of these, only the K120A mutant appears to bind as effectively as the wild-type sequence; although the binding of the D110A and R113A mutants appears to be slightly weakened, they still bind much more effectively than the F123A mutant (lane 4). These results are consistent with the hypothesis that the C terminus of MNV VPg adopts a helical conformation upon interaction with eIF4G and suggest that the helical flank in contact with eIF4G contains residues W108, D111, V115 and F123. Confirmation of this model requires further structural analysis, but if it is confirmed the helical mode of binding would likely apply to all other norovirus VPg proteins since the sequence of the eIF4G-binding motif is highly conserved (Fig 4A).

### Mapping the MNV VPg binding site on eIF4G HEAT-1

To further probe the molecular details of the MNV VPg-eIF4G interaction we used nuclear magnetic resonance (NMR) spectroscopy to map the VPg binding site on the HEAT-1 domain. ^1^H^15^N TROSY NMR experiments were used to monitor chemical shift perturbations in the spectra obtained from a solution of ^15^N labelled eIF4GI HEAT-1 domain as it was titrated with a synthetic unlabelled peptide corresponding to MNV VPg(104-124). We used the backbone resonance assignments of eIF4GI HEAT-1 deposited in the Biological Magnetic Resonance Data Bank [43] to assign changes that occurred in the ^1^H-^15^N TROSY NMR spectra on formation of the VPg(104-124):eIF4GI HEAT-1 complex to particular residues in HEAT-1.

Ideally in such experiments a limited number of amide (HN) resonances would shift (in fast or slow NMR time scales) in a dose-dependent manner. Since the spectra are dominated by signals from backbone amides (though they also contain peaks from side-chain amides in Asn, Gln, Arg and Trp residues), they can be used to map ligand binding sites. Typically clusters of surface residues that are affected by titration of the ligand are presumed to define the likely binding surface. However, titration of the VPg peptide into ^15^N labelled eIF4GI HEAT-1 was found to cause significant perturbations of most of the HN resonances, which indicates that binding of VPg affects the entire HEAT-1 domain. The exact nature of these conformational changes in HEAT-1 cannot yet be determined precisely, though they are likely to be relatively subtle since the domain retains the capacity to interact with eIF4A, which requires a large binding interface (see below and Fig 5C). As a result of the extensive perturbations, the spectrum observed at the end point of the titration (1.35 peptide:protein molar ratio) was very different to the reference spectrum obtained before the start of the titration (Fig 5A, Supplementary Fig S3). Moreover, many of the affected resonances appeared to enter a dynamics regime dominated by conformational exchange on slow or intermediate time scales, since HN peaks disappeared from the HSQC as the titration proceeded. While some new peaks appeared later in the titration, it was not possible to relate these to the peaks that had disappeared earlier in the titration (Supplementary Fig S3).

However, from closer inspection of HSQC spectra it is clear that a limited number of peaks are selectively broadened very early in the titration (Fig 5A). Since these changes happen at low peptide concentrations, they probably indicate specific contacts made by the viral protein on the eIF4GI HEAT-1 domain. To obtain a quantitative understanding of how specific HN resonances were changing early in the titration, the volume of non-overlapped peaks in the HSQCs were determined for both the reference spectrum (eIF4GI HEAT-1 only) and the spectrum obtained after addition of 0.35 molar equivalents of peptide. Volume ratios of matching peaks in the two spectra were calculated for each assigned residue (R_v_ = | Reference peak vol|)/(|0.35 eqv peak vol|) (Fig 5B). Since the number of scans recorded for the reference and for the 0.35 molar equivalent HSQC spectra were 16 and 28 respectively and the total volumes are not significantly different, R_v_ should be 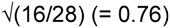 for unperturbed residues. The only HN resonances for which this ratio is approximately 0.76 are at the N- and C-termini of the domain (residues 749-973 and 977-993) and within the loop formed by residues 869-880 of the molecule, which is in a disordered region in the eIF4GII HEAT-I crystal structure [40]; these features appear not to interact with the MNV VPg C-terminal peptide. Most other residues have higher values of R_v_, as expected from the large number of perturbations caused by peptide binding. However, a small number of resonances deviate substantially from this expected value. In particular HN resonances from residues 807, 823, 825, 826, 851, 896-898, 904, 912, 914, 917-921, 935, 939 and 950 all have ratios that are more than 6 times greater than the expected value.

The position of these residues in a homology model of the eIF4GI HEAT-1 domain is given in Fig 5C. Several of these residues (823, 917, 920, and 921) are internal and probably suffer chemical shift perturbations because of conformational adjustments induced by binding of MNV VPg. Most of the other residues with very large R_v_ values cluster in two surface patches in the middle of the eIF4GI HEAT-1 domain, one formed by residues L897 and L939, which are close to the binding site for eIF4A, and one containing residues L912, E914, H918 and D919, which are on the opposite face of HEAT-1. Clearly, surface residues are more likely to be able to make direct contact with the bound MNV VPg peptide.

To confirm whether resonance perturbations assigned to residues that cluster in the surface patches in the middle of the HEAT-1 domain are involved in direct contact with MNV VPg, we tested the effect of point mutants on binding of the viral protein. The following mutations were made in eIF4GI HEAT-1: L897A and L939A in the cluster proximal to the eIF4A binding site, and L912A, H918A, D919R, and L939A in the distal cluster. (A fifth mutation in the distal cluster, E914R, was also prepared but found to have picked up an additional mutation – presumably due to a PCR error – resulting in the substitution K901M which is adjacent to the proximal cluster; this E914R/K109M double mutant therefore changes both putative VPg-binding sites). As a negative control we also made the eIF4GI HEAT-1 (K771A) mutant since this residue is in the N terminus of the HEAT-1 domain and was found in our NMR analysis to be unaffected by binding of MNV VPg (R_v_ = 0.55).

**Figure 5.**
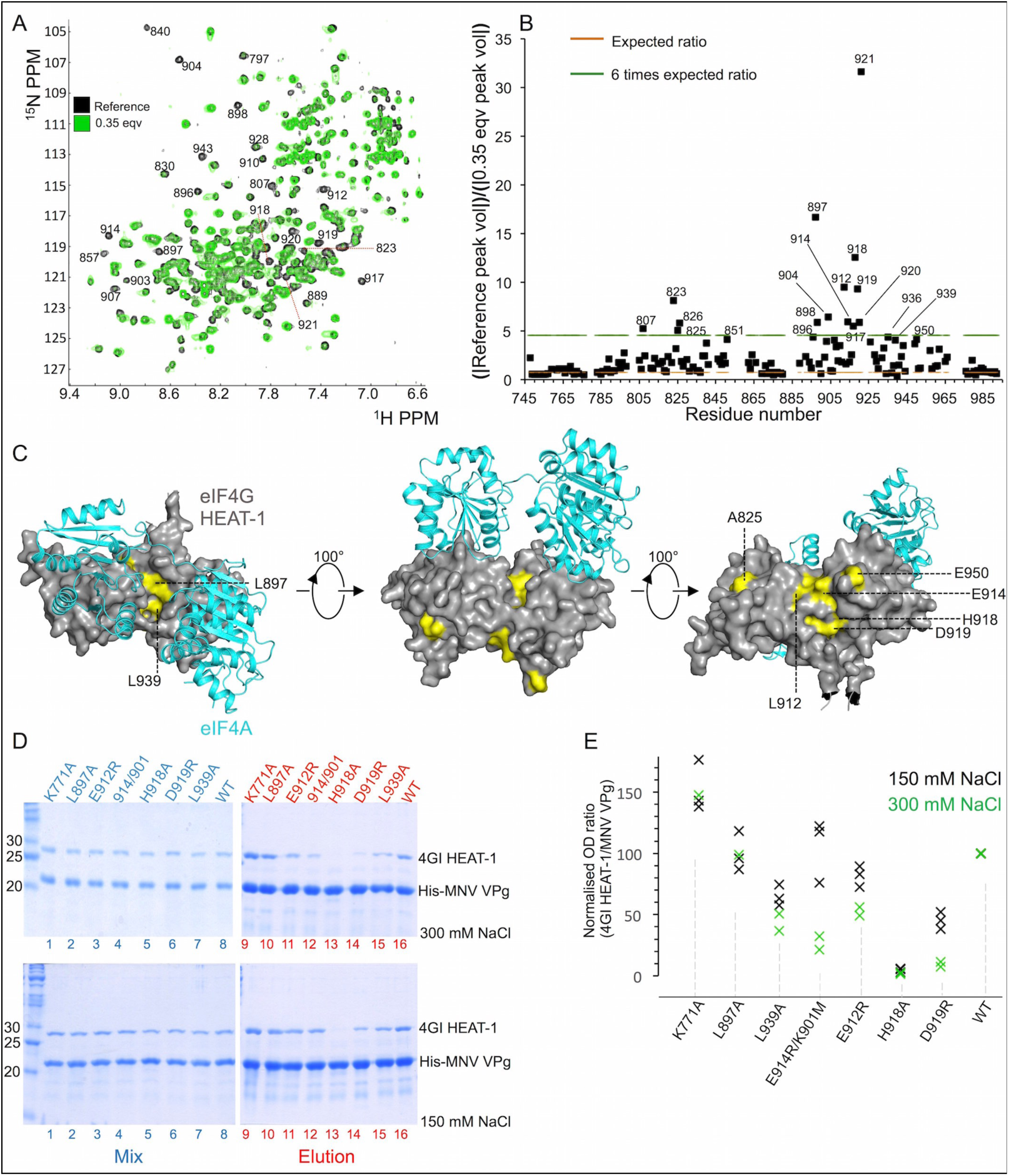
**Figure 5: NMR chemical shift mapping reveals the binding site for MNV VPg on the eIF4GI HEAT-1 domain.** (A) Superposition of the 1H15N TROSY HSQC spectra of 15N-labelled eIF4GI HEAT-1 (748-993) obtained in the absence (black) and presence (green) of 0.35 molar equivalents of unlabelled MNV VPg(104-124) peptide. (1H15N TROSY NMR spectra for all of the points in the titration are given in Supplementary Figure 3). The residues for which the intensity of amide 1H15N signals exhibit the greatest reductions as the MNV VPg(104-124) peptide concentration is increased are indicated by their residue number. The assignments were taken from the Bio Magnetic Resonance Database (BMRB id 18738). (B) Plot of the ratio of the absolute values of non-overlapping peak volumes for assigned residues obtained at 0 and 0.35 molar equivalents of the MNV VPg(104-124) peptide. High values of the ratios indicate the residues in eIF4GI HEAT-1 most affected by MNV VPg binding. The red dashed line indicates the expected volume ratio (R_v_) for unperturbed residues was 0.76 based on the relative number of scans performed in each HSQC experiment. Points above the green dashed line indicate residues for which the peak volume ratio was at least six times higher than the baseline for unperturbed amides. (C) Model of the complex of eIF4GI HEAT-1 (grey, surface representation) and eIF4A (cyan, cartoon representation) indicating the predicted location the VPg binding. Surface residues in eIF4GI HEAT-1 that exhibit the greatest changes in R_v_ in the presence of the MNV VPg(104-124) peptide are coloured yellow. The model of eIF4GI HEAT-1 was generated using SWISS-MODEL [44]; the complex was created by superposing this model on the eIF4G HEAT-1 component of the yeast eIF4G-eIF4A co-crystal structure [45]. (D) SDS PAGE analysis of pull-down assays to test the effect of mutations in the putative VPg-binding site on eIF4G HEAT-1 on binding to the viral protein. His-tagged MNV VPg(1-124) was used as bait; eIF4GI HEAT-1 wild-type or mutant proteins were used as prey, and the pull-down buffer contained 150 mM NaCl (top panel) or 300 mM NaCl (bottom panel). Lanes 1-8 (blue labels): input protein mixtures; lanes 9-16 (red labels): eluted proteins. (E) Graphical representation of the results of the cobalt affinity pull-down assays performed in the presence of 150 or 300 mM NaCl, plotted as the ratio of the optical densities of the eIF4GI HEAT-1 band to that of the His-MNV VPg band in eluted fractions (lanes 9-16 in panel D). Band densities were quantified using ImageJ (http://imagej.nih.gov/ij/) and the ratios were normalised to the wild-type control.

The VPg-binding ability of these eIF4G mutants was tested in a cobalt affinity pull-down assay in which His-MNV VPg(1-124) was used as bait. A first series of assays was performed using a binding buffer containing 150 mM NaCl (see Materials and Methods). This demonstrated that the H918A mutation in the distal cluster had the most severe defect in MNV VPg binding, reducing it to less than 10% of the control, while nearby substitutions D919R and L939A showed more modest decreases in binding (Fig 5D-E). Within the proximal cluster, the L897A mutation had no effect on binding, while the L939A mutation reduced binding to about 60% of the control value. To increase sensitivity, a second series of assays was performed in buffer containing 300 mM NaCl. Under these higher-salt conditions, binding defects due to the mutations were more pronounced. The H918A mutant was again found to exhibit the weakest binding of MNV VPg, but the deleterious effects of the E912R and D919R mutations were more severe. In the proximal cluster, there was still no effect of the L897A mutation but the binding defect due to the L939A mutation was slightly worse. The interaction of VPg with the E914R/K901M double mutant, which was similar to wild-type at 150 mM NaCl, was reduced significantly at a salt concentration of 300 mM. Finally, as expected, the negative control mutation K771A had no effect on binding at low or high salt (Fig 5D-E).

These results are consistent the location of the MNV VPg binding site on eIF4GI HEAT-1 that was mapped in NMR titration experiments. They also suggest that the binding site is centred on the surface patch (around residues 914-919) distal to the eIF4A binding site. The binding site may extend to residue L939 but residue L897, which did not affect binding of VPg when mutated and is closest to the binding site of eIF4A, may define an upper boundary of the VPg binding site. It is perhaps worth noting that if residues 108-124 of MNV VPg were to form a helix on binding to eIF4G (see above), the helix would have five turns and therefore be of a similar length to the helices in the HEAT-1 domain.

### MNV VPg forms a stable complex with eIF4A and eIF4G HEAT-1 at a stoichiometry of 1:1:1

The VPg-binding site on eIF4GI identified by our NMR and mutagenesis experiments lies on a surface of the HEAT-1 domain that is distinct from the eIF4A-binding site. This suggested to us that VPg binding would not interfere with the eIF4G-eIF4A and that MNV VPg might therefore be able to form a ternary complex with eIF4G and eIF4A. The presence of eIF4A in initiation complexes formed with MNV VPg is consistent with earlier observations that MNV VPg translation is sensitive to inhibition of the ATPase activity of eIF4A by hippuristanol [27].

We used size exclusion chromatography (SEC) to characterise the complexes formed by MNV VPg and eIF4A with eIF4GI HEAT-1. Stoichiometric mixes of MNV VPg and eIF4GI HEAT-1, and eIF4A and eIF4GI HEAT-1 both resulted in peaks that eluted at lower volumes (14.7 and 14.3 mL respectively) than observed for the individual component proteins, consistent with the formation of binary complexes (Fig 6A). The presence of these complexes was confirmed by SDS PAGE analysis (Fig 6A inset). An equimolar mix of 1:1:1 MNV VPg, eIF4GI HEAT-1, and eIF4A eluted at an even lower volume (13.6 mL), consistent with the formation of a complex involving all three proteins, an interpretation that was also confirmed by SDS PAGE analysis (Fig 6A).

The SEC analysis was coupled to multi-angle laser light scattering (SEC-MALLS) to measure the molecular weights (MW) of these complexes and determine their stoichiometry (Fig 6B). For the VPg:eIF4G HEAT-1 complex the observed MW (45.9 kDa) was very close to that expected for a 1:1 complex (46.5 kDa). For the eIF4A:eIF4G HEAT-1 complex the observed 65.9 kDa MW was somewhat lower than the 79.7 kDa expected for a 1:1 complex. The reason for this small discrepancy is not clear – it may be due to partial dissociation during elution – but the formation of a 1:1 binary complex is still the most likely outcome. The average MW measured for the ternary MNV VPg:eIF4A:eIF4GI HEAT-1 complex using SEC-MALLS was 94.5 kDa, which is very close to the 94.1 kDa expected for a complex of 1:1:1 stoichiometry. The formation of this ternary complex is consistent with the observation that MNV VPg and eIF4A have non-overlapping binding sites on the eIF4G HEAT-1 domain. This observation also helps to explain how MNV VPg is able to interact with functional eIF4E:eIF4G:eIF4A complexes, as shown by the ability of the protein to interact with these proteins in cell lysates [39]; the eIF4E binding site on eIF4G is upstream of the HEAT-1 domain within eIF4G [9], allowing eIF4E to bind independently of eIF4A and MNV VPg.

**Figure 6.**
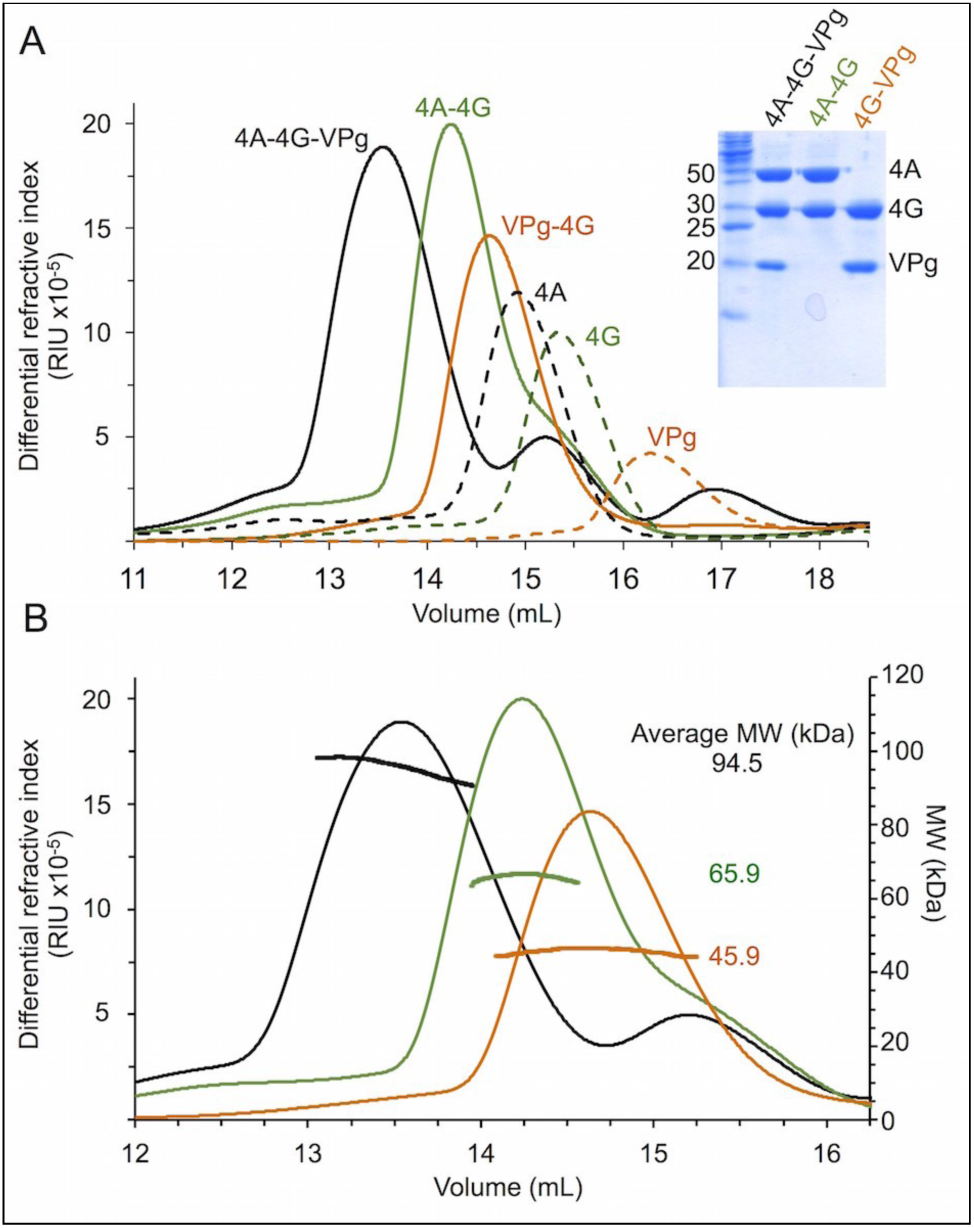
**Figure 6: MNV VPg forms a ternary complex with eIF4G and eIF4A.** (A) SEC profiles monitored by differential refractive index (which is proportional to protein concentration) for eIF4A, eIF4G HEAT-1 and MNV VPg as well as for the binary complexes formed by mixing approximately equimolar quantities of eIF4A and eIF4G HEAT-1 (4A-4G), eIF4G HEAT-1 and MNV VPg (4G-VPg), and the ternary complex obtained from eIF4A, eIF4G HEAT-1, and MNV VPg (4A-4G-VPg). SDS PAGE analysis of peak fractions of the binary and ternary complexes obtained in the SEC experiments are also shown in A. (B) SEC-MALLS analysis of the molar mass distributions of the binary and ternary complexes plotted against the SEC profiles shown in A.

### MNV VPg C-terminal peptide fusions inhibited MNV VPg mediated translation *in vitro* but could not be shown to have any anti-viral activity

Our *in vitro* binding studies have shown that a conserved motif within the C terminus of MNV VPg mediate a micromolar affinity interaction with the HEAT-1 domain of eIF4GI and eIF4GII, possibly by adopting a helical conformation that interacts with a centrally-located binding site that does not overlap with the binding site for eIF4A. To probe the physiological significance of this observation we tested whether GST-fusions containing the C terminus of MNV VPg could inhibit norovirus translation. To do so we performed *in vitro* translation assays programmed with VPg-linked genomic RNA in the presence of increasing concentrations of GST-MNV VPg(102-124) or the GST-MNV VPg(102-124) F123A mutant, which binds eIF4G much more weakly. As shown in Fig 7A GST-MNV VPg(102-124) inhibits norovirus translation in a dose-dependent manner. At the highest concentration used (28 µM) the yield of translation product is reduced to 40% of the control value. In contrast, at all concentrations the non-binding GST-MNV VPg (102-124) F123A mutant causes no significant inhibition on translation.

**Figure 7.**
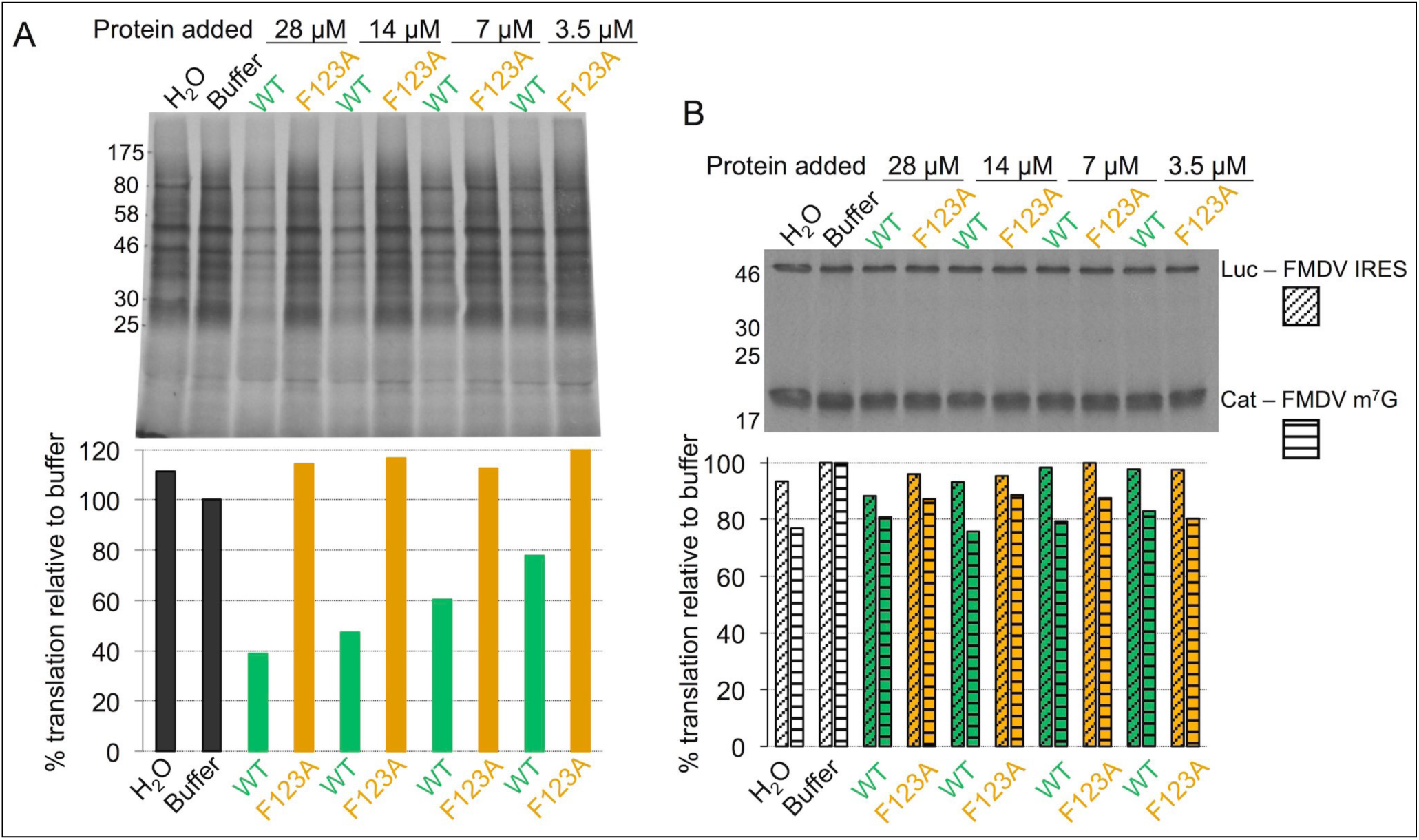
**Figure 7: GST-MNV VPg 102-124 inhibits MNV VPg-mediated translation *in vitro* but not cap-dependent or IRES-dependent translation.** *In vitro* translation reactions were performed in the presence of increasing concentrations of GST-MNV VPg(102-124) WT protein or the GST-MNV VPg(102-124) F123A mutant that binds much less well eIF4G. Protein synthesis was monitored by autoradiography of SDS PAGE analysis of incorporation of 35S-methionine in translation reactions. (A) Top panel: Effect of exogenous GST-MNV VPg(102-124) proteins on translation from VPg-linked MNV RNA; bottom panel: quantitative analysis of the level of 35S-methionine incorporation. (B) Top panel: Effect of exogenous GST-MNV VPg(102-124) proteins on translation from capped bi-cistronic mRNA constructs containing the FMDV IRES between the first (CAT) and second (Luc) cistrons; bottom panel: quantitative analysis of the level of 35S-methionine incorporation.

As a further test of the specificity of the inhibition of VPg-mediated translation the same proteins were used at the same concentrations in alterative *in vitro* translation assays. These were programmed with a capped bicistronic construct containing an open reading frame (ORF) for chloramphenicol acetyl transferase (CAT) under the control of an m7G cap and a downstream luciferase (LUC) ORF under the control of an IRES. Two different constructs were tested, one with a FMDV IRES and the other with a PTV IRES. We found that neither the wild-type nor the F123A mutant form of GST MNV VPg (102-124) had any effect on translation of either CAT or LUC at even the highest protein concentrations, irrespective of the type of IRES present between the cistrons (Fig 7B, Supplementary Fig S4). This suggests that the inhibition of VPg-dependent translation by exogenous added GST-MNV VPg(102-124) is specific. It is likely mediated by competition for binding to eIF4G.

As a second probe of the physiological relevance of our observation that the C terminus of MNV VPg mediates binding of the protein to eIF4G, we tested whether MNV VPg(102-124) could interact with the large ribosomal initiation complexes in eukaryotic cells, as has been previously demonstrated for TAP-tagged MNV VPg(1-124) [39]. To do this plasmids encoding mCherry-MNV VPg(102-124) and mCherry-MNV VPg(102-124) F123A fusion proteins were expressed in BV2 macrophages using lentivirus vectors (see Materials and Methods). Cell lines expressing mCherry MNV VPg(102-124) wild-type or the F123A mutant were selected on the basis of equal fluorescence intensity. Subsequent α-mCherry co-immunoprecipitation and western blot analysis revealed that eIF4G, eIF4A, eIF4E, eIF3D and PABP all co-immunoprecipitate with wild-type mCherry MNV VPg(102-124) but not with the F123A version (Fig 8A). This suggests that the MNV VPg C-terminal peptide can form the same complexes in the cytosol as the full-length protein.

Finally, since we had shown that MNV VPg(102-124) can inhibit noroviral translation *in vitro* and interacts specifically with initiation complexes in cells, we sought to determine whether cells over-expressing mCherry MNV VPg(102-124) are resistant to infection by the virus. However, no difference was observed in MNV titre over a 24-hour time course between cells expressing mCherry MNV VPg (102-124), or the non-binding mCherry-MNV VPg (102-124) F123A mutant or mCherry alone (Fig 8B).

**Figure 8.**
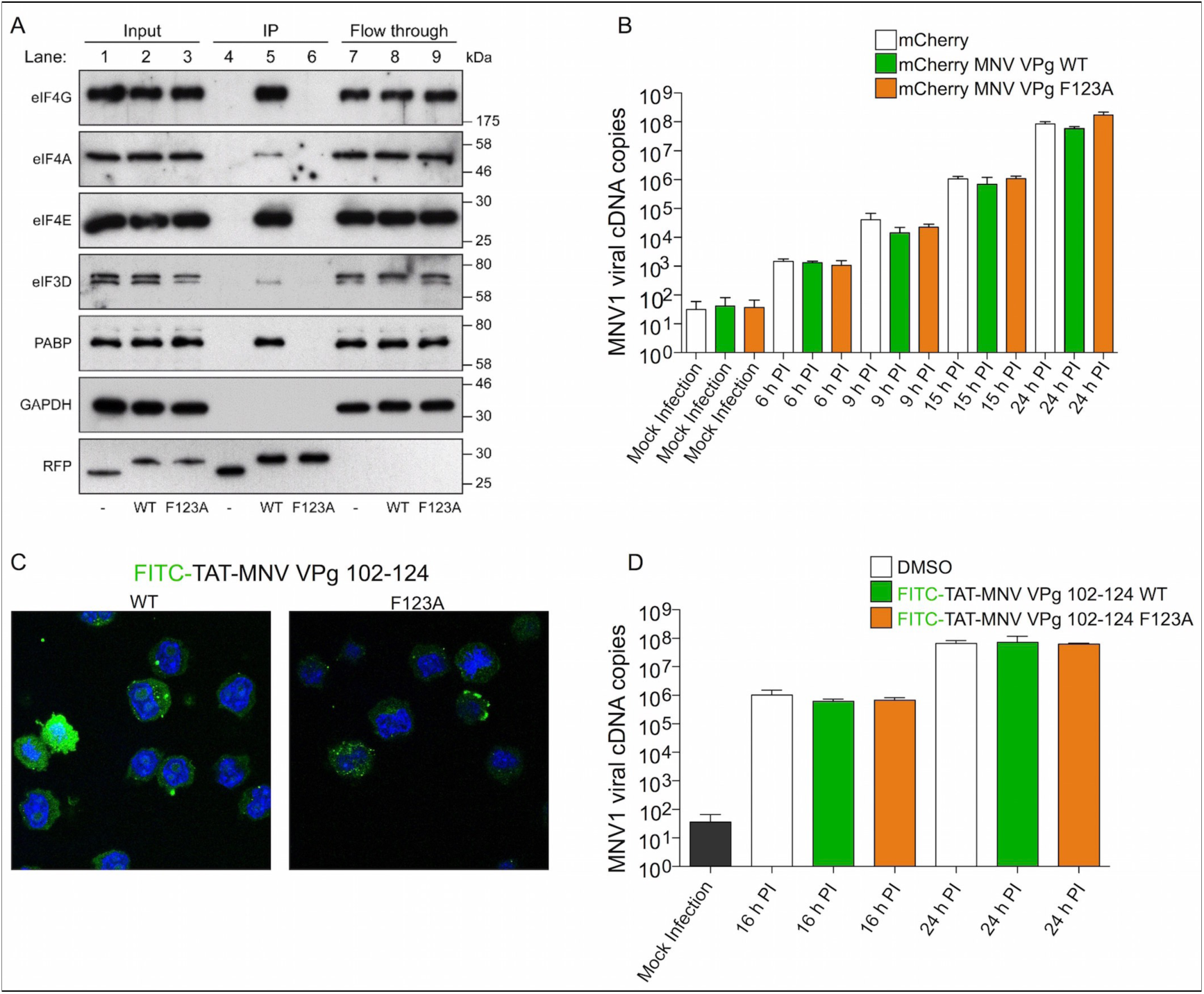
**Figure 8: mCherry-MNV VPg(102-124) co-immunoprecipitates with translation initiation proteins.** (A) RFP-Trap(r) immunoprecpitation of lysates of BV2 cells expressing mCherry, mCherry-MNV VPg(102-124) WT or mCherry-MNV VPg(102-124) F123A. Pre-purification lysates (input), purified fractions (IP) and unbound fractions (Flow through) were analysed by SDS PAGE and western blotting. (B) Time course of BV2 infection (MOI 0.01 TCID_50_ units/cell) with MNV1 in the presence of mCherry-VPg proteins. Prior to infection BV2 cells were transduced with lentiviruses expressing mCherry, mCherry-MNV VPg(102-124) WT or mCherry-MNV VPg(102-124) F123A. The progress of infection was monitored by measuring the number of viral cDNA copies generated from quantitative RT-PCR analysis of whole-cell RNA. (C) Fluorescence microscopy analysis of cell penetration of the FITC-TAT-MNV VPg(102-124) peptides. The images shown are merged images of DAPI stained nuclear DNA (blue) and wild-type or F123A versions of the cell penetrating peptides (green). (D) Time course of BV2 infection (MOI 0.01 TCID_50_ units/cell) with MNV1 following pre-treatment for 150 minutes with 100 µM of wild-type or F123A versions of the cell penetrating peptides prior to infection. The progress of infection was monitored by RT-PCR analysis (as in panel B).

The mCherry tag adds more than 25 kDa to the molecular weight of the peptide. The lack of inhibition of norovirus replication might therefore be due to the copy number of the molecule in cells being too low. As an alternative test of the inhibitory activity of the MNV VPg C terminus wild-type and F123A variants of the MNV VPg(102-124) that incorporated N-terminal cell penetrating peptide (CPP) sequences were synthesised. Two different CPP sequences were used, derived from HIV TAT and Bac7. In each case the peptides were also N-terminally tagged with FITC so that penetration into cells could be monitored with fluorescence microscopy. As shown in Fig 8C and Supplementary Fig S5A, the peptides were able to get into the cytoplasm. However, no difference was observed in viral titre over a 24-hour time course between cells containing either wild-type or F123A versions of the cell penetrating VPg peptides when compared to controls that had only been exposed to the peptide solvent, DMSO (Fig 8D and Supplementary Fig S5B). The reason for the negative result is not clear although it may again be due to delivery of insufficient peptide to the cell interior.

## Discussion

It has been known for nearly 20 years that the VPg protein covalently attached to the 5′-end of calicivirus RNA genomes is critical for translation [28]. Since then, various interactions of VPg proteins with eukaryotic initiation factors, including eIF3, eIF4E and eIF4G, have been reported [27,35-37,39]. Of these, the interaction of FCV VPg with eIF4E and of MNV VPg with eIF4G appear to be the most functionally significant [27,39] but mechanistic details beyond the idea that VPg serves to recruit the viral RNA to the translation machinery have been lacking.

In this paper we report the discovery and characterisation of a conserved ∼20 amino acid motif in the C terminus of norovirus VPg proteins that binds to the HEAT-1 domains of eIF4GI and eIF4GII with micromolar affinity, an interaction that is crucial for the initiation of translation from the viral RNA genome. Norovirus VPg therefore serves as a proteinaceous ‘5′-cap’ that does not require a functional interaction with the cap-binding protein, eIF4E [27]. Our data provide clear evidence for the mechanistic basis for an important and unusual paradigm of translation initiation that is independent of 5′-cap and IRES structures, and place functional and geometric constraints on how it operates. We have shown that the equilibrium dissociation constant for binding of the C terminus of VPg for the eIF4G HEAT-1 domains is in the low micromolar range – around 3 µM (Fig 3). This is relatively weak when compared to the affinity of m7G capped mRNA for the eIF4F complex which is in the low nanomolar range [46]. IRES RNA structures are also reported to bind to initiation factor complexes with affinities in the low nanomolar range [47-49]. At first sight it may therefore appear questionable whether the binding of MNV VPg to eIF4G HEAT1 is of high enough affinity to recruit the ribosome, particularly if it is in competition with 5′-capped eukaryotic mRNAs. However, it seems likely that in infected cells the VPg:HEAT-1 interaction forms just one contact in the overall interaction between VPg-linked genomic RNA and the initiation machinery, though we suspect that these points of contact remain focused on eIF4G. Although initial interaction of the viral RNAs with eIF4F leads to recruitment of eIF3 and the 40S ribosomal subunit, and there is evidence that norovirus VPg can bind to both eIF4E [37] and eIF3 [35,36], the functional significance of these interactions has yet to be demonstrated. Indeed, although there is residual binding of MNV VPg to eIF4E when the eIF4G binding site on the viral protein has been mutated, abrogation of eIF4E-dependent translation initiation, either through the addition of the 4E-binding protein or cleavage of eIF4G by the FMDV L protease, has no effect on translation of MNV VPg-linked RNA [27].

It seems more likely that RNA interactions play a significant role in the formation of initiation complexes with VPg-RNA, since capped and uncapped mRNA can both bind to eIF4F with nanomolar affinity [46], mostly likely though interactions with RNA binding sites on eIF4G and eIF4A [40]. Although the cap adds little to the overall affinity, it may serve to specify the orientation of cellular mRNA binding to eIF4F since the interaction with RNA is not otherwise sequence-specific. It may be that VPg is acting in a similar mechanism to the m7G cap, providing an important topological constraint for a high-affinity interaction between noroviral RNA and the eIF4F complex to ensure efficient initiation.

The precise geometry of the complex formed when noroviral RNA binds to eIF4F is difficult to determine. Although there is a crystal structure for the binary complex comprising eIF4A and the HEAT-1 domain of eIF4G [45] and we have at least partially mapped the binding site on eIF4G HEAT-1 for the conserved C-terminal peptide in MNV VPg (Fig 5), the C terminus of VPg is linked flexibly to the helical core of the protein (residues 11-62) which contains the covalent point of attachment (Tyr 26) of the viral RNA [34]. Since the AUG start codon in noroviral RNA is only 4 nucleotides from the 5′-end, the length of this linker may be required to permit positioning of the start codon in the ribosomal P-site without it being encumbered by the bulk of the eIF4F complex. Similar considerations apply to all noroviruses since the key functional features – proximity of the start codon to the 5′-end of the viral RNA, site of RNA attachment to VPg and length of the linker to the conserved eIF4G-binding motif – are conserved. However, high-resolution structural analyses will be required to establish the relative positions of norovirus VPg-RNA and eIF4F within the context of the 48S ribosomal initiation complex.

Similar geometric constraints are likely to affect the translation initiation mechanisms of other genera within the *Caliciviridae*, even though their VPg proteins lack the eIF4G-binding motif found in norovirus VPg. For example, the best available evidence suggests that binding of FCV VPg to eIF4E is required for translation initiation [27,37]. This positions the FCV VPg at a different locus within the eIF4F complex compared to MNV VPg, though their binding sites on eIF4F may not be far apart – the eIF4E binding site in eIF4G (residues 557-681) is close to the HEAT-1 domain (residues 751-1011) [9]. Intriguingly, the mechanism reported here for noroviruses may also operate in rice yellow mottle virus (RYMV) since there is evidence to suggest that its VPg protein binds to the eIF4G HEAT-1 domain [50] and appears to have a C-terminal sequence that is similar to the HEAT-1 binding motif identified in norovirus VPg (Fig 4A).

The observation that the VPg-NS6pro fusion protein cannot bind to the eIF4G HEAT-1 domain (Fig 2D) shows that the C terminus of VPg has to be free for a productive interaction, and raises interesting questions about the regulation of norovirus replication. As with all positive-sense, single-strand RNA viruses, the early stages of norovirus infection are dominated by translation, which allows the accumulation of copies of the polyprotein precursor. This precursor is eventually processed by NS6pro into the functional proteins required for virus replication. The precise timings of the processing cascade are not known in detail, although evidence suggests that cleavage of VPg-NS6pro is significantly slower than the early cleavages at the NS2-3 and NS3-4 sites [51,52]. It may well be that the timing of the processing of precursors containing VPg is tuned to permit an early burst of translation (unimpeded as long as VPg remains fused to NS6^pro^), followed by a switch to RNA replication that may be facilitated by the accumulation of free VPg (or precursors with VPg at the C terminus), which would bind to eIF4G and inhibit viral translation initiation. This might help to clear the viral RNA of ribosomes to facilitate initiation of RNA replication, which requires association of the viral NS7^pol^ at the 3′-end followed by movement along the RNA template in the 3′-5′ direction. Consistent with this notion, we have shown that exogenous GST-VPg fusion proteins containing the eIF4G-binding peptide can inhibit translation from norovirus VPg-RNA (Fig 7A), but it remains to be seen if this hypothetical mechanism plays out in infected cells.

Finally the discovery of a binding site on eIF4G that is critical for norovirus translation initiation raises the possibility that the VPg-eIF4G interaction might be targeted in the development of antiviral compounds. Our initial efforts to demonstrate the antiviral of VPg peptides failed to show any effect (Fig 8, Supplementary Fig S5), but this could simply be because an insufficient cytoplasmic concentration was achieved. In any case, small molecule inhibitors are likely to have improved pharmacokinetics and our results lay the necessary groundwork to enable high-throughput screening for compounds that might disrupt the VPg-eIF4G interaction and so block infection.

## Materials and Methods

### Plasmids for protein expression in *E. coli.*

Six different expression vectors were used to make new expression constructs. Briefly these were pETM-11 which has been described elsewhere [53]. Two modified versions of pETM-11, one with a *Bam*HI site replacing the *Nco*I site (pETM-11m1), and the other, a variant of pETM-11m1, in which a thrombin cleavage site downstream of the N-terminal His-tag replaces the Tobacco Etch Virus (TEV) NIa cleavage site (pETM-11m2) [54,55]. pMALX(E) has been described elsewhere [56] and pGEX-2T is commercially available (GE healthcare). An additional modified pGEX-2T plasmid was used with a TEV NIa cleavage site replacing the thrombin site (pGEX-2Tm). eIF4GI, eIF4GII and DAP5 sequences were amplified by PCR (KOD polymerase hot start kit) from full length cDNA clones.

cDNA for eIF4GI, eIF4GII and DAP5 were kindly supplied by Chris Hellen and Tatyana Pestova (State University New York), Mark Coldwell (University of Southampton) and Bhushan Nagar (McGill University, Montréal) respectively. The residue numbering used throughout this paper for these proteins is based on the following NCBI accession codes, eIF4GI (AAM69365.1), eIF4GII (NP_003751.2) and DAP5 (NP_001036024.3). The oligonucleotide primers used in amplification of cDNA are given in Supplementary Table S1. Ligation into expression vectors was performed using the T4 Quick Ligation Kit (New England Biolabs).

Fusions of glutathione-S-transferase (GST) with VPg C-terminal peptide sequences were made by ligating annealed synthetic oligonucleotides into pGEX-2T (Supplementary Table S1). 5 µM of sense and anti-sense oligonucleotide were heated to 95°C for 12 minutes in NEB buffer 2 followed by slow cooling to room temperature over 2.5 hours. The oligonucleotides were designed so that annealing would generate sticky ends corresponding to the *Bam*HI and *Eco*RI restriction sites that were used in ligation.

Green-fluorescent protein (GFP) fusions were made in the following way: cDNA for enhanced GFP-VPg and GFP-VPg|NS6 fusions were generated by using overlap extension PCR to fuse enhanced GFP to MNV VPg 1-124 using a <u>ENLYFQG</u>AAA linker between the two proteins that incorporates contains a TEV NIa Cleavage site (underlined). The PCR product was ligated into pETM-11m1.

Full length hexa-Histidine-tagged VPg constructs (MNV VPg(1-124) wild-type (WT) and the F123A mutant, and FCV VPg(1-111) WT), and GST-eIF4GI 4GM (751-1132) constructs are described elsewhere [34,39]. pET15b-eIF4A was kindly supplied by Chris Hellen and Tatyana Pestova (State University New York) [57,58].

Almost all mutations were made by QuikChange site directed mutagenesis (Stratagene). The exception to this was the MNV VPg(102-124) K120A mutant, which was made by annealing oligonucleotides and ligating the product as described above. Details of all of the constructs used in the study, including residue numbers, expression vector, purification tag details, protease cleavage site to remove the tag and non-native residues in the sequence post-cleavage are summarised in Supplementary Table S2. All constructs were verified by DNA sequencing (MWG Operon).

### Plasmids for protein expression in BV2 cells

Lentivirus vectors expressing mCherry-MNV1-VPg fusions were generated in two steps using the plasmid pTM900, a bicistronic lentiviral vector derived from pCCLsin.PPT.hPGK.GFP.pre that carries a multiple cloning site downstream of the human phosphoglycerate kinase promoter and coexpresses a hygromycin resistance gene under the control of the EMCV IRES [59,60]. The mCherry coding sequence lacking an in-frame stop codon was first amplified by PCR (see Supplementary Table 2 for primer details) and cloned into the lentiviral vector pTM900 between the *Bsh*TI and *Bsi*WI restriction to replace the open reading frame for green fluorescent protein (GFP). In the second step, the coding sequence of wild-type and mutated VPg peptides were inserted in frame with the mCherry ORF by hybridizing complementary single-stranded oligonucleotides encoding the last 23 amino acids of MNV1 VPg flanked by *BsiW*I and *Xba*I restriction sites and ligated into the corresponding sites of the vector generated in the first step. The sequences of the oligonucleotides used to amplify MNV VPg wild-type and F123A mutants for cloning into the lentivirus vector are given in Supplementary Table S2. A bicistronic lentiviral vector co-expressing mCherry and a hygromycin resistance gene was used as negative control. All plasmid sequences were verified by DNA sequencing.

### Protein expression and purification

All proteins were expressed in *E. coli* BL21-CodonPlus(DE3) RIPL or BL21-Rosetta(DE3) cells. The cells type used for each construct is given in Supplementary Table S3. For non-isotopically labelled proteins 1 L of lysogeny broth (LB) was seeded using 50 mL of an overnight starter culture. The cells were grown at 220 RPM and 37°C until mid-log phase was reached (OD_600_ of ∼0.6-0.8) at which point the cells were typically induced with 1 mM isopropyl β-D-1-thiogalactopyranoside (IPTG) and allowed to grow at 220 RPM and a defined temperature and period of time. Cells were pelleted by centrifugation at ∼5,000 g and stored at -80°C. The temperature and time regimen used for each protein are listed in Supplementary Table S3.

For expression of 15N labelled eIF4GI HEAT-1 four 1 L cultures of LB were seeded and grown to mid-log phase. At this point the cells were pelleted at ∼3,000 g for 20 minutes. The supernatant was discarded and the cells re-suspended in 800 mL of sterile TBS (50 mM Tris pH 7.5, 150 mM NaCl). The centrifugation step was repeated, the supernatant discarded and the pellets finally re-suspended in a total of 1 L of minimal medium divided between two 1 L culture flasks. The minimal medium, which contained ^15^NH_4_Cl as a labelled nitrogen source, was prepared as detailed in Supplementary Table S4. Protein expression was induced after 2 hours at 30°C by addition of 1 mM final IPTG and continued for approximately 16 hours at 30°C and 220 RPM. The cells were harvested and stored as previously described.

The purification of the various recombinant proteins from *E. coli* followed the general scheme presented below. The details of the buffers used for each protein purification are fully documented in Supplementary Table S5 (and cross-referenced to the data presented in each figure); any significant deviations from the protocol below are given in the supplementary methods.

Cells were re-suspended in a purification buffer (spiked with 2 mg/mL final Chicken egg lysozyme, 0.1 mM phenylmethanesulfonyl fluoride (PMSF) and 0.1% Triton X-100) and lysed by sonication in 15-second pulses separated by 15-second rests on ice over a period of 5 minutes. Lysates were clarified by centrifugation at 29,000 g for 20 minutes. DNA was precipitated by addition of protamine sulfate to a concentration of 1 mg/mL and the lysate centrifuged again at 29,000 g for 20 minutes. Clarified lysates were applied to their respective affinity resins: TALON (Clontech) or His-select (Sigma) for His-tagged protein, Glutathione 4B (GE Healthcare) resin for GST tagged proteins, or amylose resin (New England Biolabs) for MBP-tagged proteins. In each case the lysate was slowly rotated with the resin for ∼1 h at 4°C and then applied to a gravity flow column. The washing and elution strategy employed was dependent on the affinity strategy being employed. For His-tagged proteins the resin was washed sequentially with 25 mL batches of purification buffer with increasing concentrations of imidazole. Proteins typically eluted at an imidazole concentration of 100 mM. For GST-tagged proteins the resin was washed with four 25 mL batches of purification buffer and eluted with two washes of the same buffer containing 10 mM reduced glutathione, the first titrated to pH 8 the second to pH 9; MBPtagged proteins bound to the amylose resin were washed in the same way but the protein was eluted using 15 mL of purification buffer containing 10 mM maltose. Pure fractions were dialysed overnight (14-17 hours) against 4 L of dialysis buffer (Supplementary Table S5). For proteins in which the affinity tag was removed this was done by addition of protease (typically 0.5 mg TEV NIa or 200 U Bovine thrombin for 2 L cultures) at the beginning of dialysis.

Following dialysis, proteins which had not been processed to remove tags were concentrated using centrifugal concentrator, snap frozen in liquid nitrogen and stored at 80°C. Proteins that were proteolytically-processed during overnight dialysis were re-applied to the appropriate affinity resin to remove the cleaved tag. The flow-through fractions obtained at this stage, which contained the untagged protein, were concentrated in a centrifugal concentrator and further purified by size exclusion chromatography (SEC), typically using a Superdex 75 16/60 column connected to an ÄKTA FPLC (GE Healthcare). The SEC buffers are detailed in Supplementary Table S5. Pure fractions were pooled and concentrated using centrifugal concentrator and snap frozen in liquid N_2_ and stored at -80°C. Protein concentrations were determined from optical density measurements at 280 nm using calculated extinction coefficients; peptide concentrations were calculated from the dry weight of synthesised material.

### Glutathione affinity pull-down assays

These assays were performed to probe the binding of various glutathione-S-transferase (GST) tagged eIF4G proteins (used as bait) to N-terminally His-tagged MNV VPg (1-124) (prey). In a typical experiment, 30 µL of the GST-eIF4G and MNV VPg proteins were mixed in SigmaPrep spin columns with 430 µL of Tris binding buffer (50 mM Tris pH 7.6, 150 mM NaCl), and 75 µL bed volume of glutathione sepharose 4B resin (GE healthcare). For these assays final protein concentrations were 1.4 µM bait and 2.3 µM prey. The mixture was allowed to rotate slowly at 4°C for 60 minutes prior to collection of the flow through by centrifugation at 100 g for 30 seconds. The resin was washed 3 times with 750 µL of Tris binding buffer by centrifugation. The protein was eluted by incubating the resin for 2 minutes with 75 µL of Tris elution buffer (100 mM Tris pH 8, 300 mM NaCl, and 10 mM reduced glutathione) prior to centrifugation at 2,000 g for 2 minutes. Pre-purification mixtures and eluate samples were analysed by SDS PAGE and Coomassie staining.

### Cobalt affinity pull-down assays

These assays were usually performed using N-terminally His-tagged versions of FCV VPg 1-111 or MNV VPg 1-124 as bait and N-terminally GST-tagged eIF4G constructs as the prey protein (see individual figures for details). Before use GST-tagged prey proteins were dialysed overnight (at a volume ratio of 1/1000) into 25 mM 2-(*N*-morpholino)ethanesulfonic acid (MES) pH 6.5, 150 mM NaCl, 2 mM 2mercaptoethanol. The VPg bait proteins were in a buffer containing 15.2 mM Na_2_HPO_4_, 34.2 mM NaH_2_PO_4_, pH6.5, 300 mM NaCl and 1 mM DTT. For each assay typically a 40 µL mixture of bait and prey proteins was added to 450 µL of phosphate binding buffer (9.2 mM NaH_2_PO_4_,1.8 mM KH_2_PO_4_, 136.9 mM NaCl, 2.7 mM KCl, 1 mM 2-mercaptoethanol, which was titrated to pH 7.4 with NaOH) in a SigmaPrep spin column along with 25 µL bed volume of TALON resin (Clontech). Typical final protein concentrations were 1 µM for the bait and 6 µM for the prey (see figure legends for details). The mixture was rotated slowly at 4°C for approximately 70 minutes. Following incubation the unbound fraction was collected by centrifugation of the spin columns at 100 g for 5 seconds. The resin was then washed by centrifugation, once with 750 µL phosphate binding buffer, and then twice with 750 µL binding buffer containing 10 mM imidazole. Bound proteins were eluted by incubating the resin for 2 minutes with 50 µL of phosphate binding buffer containing 250 mM imidazole, followed by centrifugation for 2 minutes at 2,000 g. Samples of eluates and protein mixes taken before application to the resin analysed by SDS PAGE and Coomassie staining.

### Fluorescence anisotropy binding assays

Fluorescence anisotropy was used to measure the binding of synthetic VPg peptides (labelled at the N terminus with fluorescein isothiocyanate (FITC)) to HEAT-1 domains from eIF4G1, eIF4GII and DAP5. Measurements were performed at 29°C using a Spectramax i3 Spectrometer with excitation/ emission wavelengths of 485 and 535 nm respectively, slit widths set to 20 nm (excitation) and 25 nm (emission) and an integration time of 400 ms. Samples were loaded into non-binding surface black half area plates (Corning) using a MES binding buffer composed of 25 mM MES pH 6.5, 150 mM NaCl, 3 mM DDT, 0.05% Tween-20.

N-terminally FITC labelled peptides MNV VPg 104-124 (VGPSWADDDRQVDYGEKINFE-COOH) and MNV VPg 108-124 were purchased at >95% purity from ChinaPeptides Co., Ltd. (Shanghai, China). In both cases the FITC group was attached with via a 6-aminohexanoic acid group to the N terminus of the peptide. Peptides were dissolved in binding buffer and titrated to pH 6.5 using a small volume of concentrated NaOH. The final peptide concentration was kept at a constant 10 nM across all binding experiments. The proteins tested for binding – untagged eIF4GI HEAT-1 truncated (748-993), untagged eIF4GII HEAT-1 (745-1003), MBP tagged eIF4GII HEAT-1 (745-1003) and DAP5 HEAT-1 (61-323) – were all dissolved in MES binding buffer. For each experiment a two-fold dilution series of each HEAT-1 protein was performed (typically starting at 20 or 40 µM) using MES binding buffer as diluent. For each dilution series, a no-protein control was performed in which binding buffer replaced protein. Each measurement typically involved two independent dilution series performed in parallel. The average fluorescence anisotropy values for the no-protein controls were subtracted from the fluorescence anisotropy values for each point in the titration giving the difference (ΔFA) for each data point. The ΔFA values for each point in the titration were used to calculate the dissociation constant (K_D_) for each interaction tested. This was done by fitting all of the titration data to a single-site binding model (ΔFA = FA_max_^*^[protein]/(K_D_ + [protein]) where FA_max_ is the maximal change in fluorescence anisotrpy) in Prism6 (GraphPad Software).

Titration of up to 40 µM eIF4GI into an irrelevant peptide ([FITC]-GSSHRYFLERGLESATSL; [61]) did not result in significant increases in fluorescence anisotropy (ΔFA < 0.004), confirming the specificity of the interaction with the MNV VPg peptides.

### Isothermal Titration Calorimetry (ITC)

For ITC experiments MNV VPg 1-124 and eIF4GII HEAT-1 (745-1003) – both with their His-tags removed – were dialysed overnight into 1 L of MES binding buffer using Sigma-Pur-a-lyzer midi dialyser (3,500 kDa MWCO). Following dialysis the protein stock concentrations were determined by measurement of absorption at 280 nm to be 52 µM MNV VPg(1-124) and 544 µM eIF4GII HEAT-1. ITC experiments were performed at 29°C using a MicroCal iTC200 instrument by injecting small volumes of eIF4GII HEAT-1 in 200 µL MNV VPg(1-124) in the sample cell. A total of 20 injections were performed, the first of 0.5 µL followed by 19 injections of 2 µL. The first two injections were separated by 180 s; all subsequent injections were separated by 120 s. Integrated raw data analysis was performed using Origin9 software using the single-site binding model to determine the values of K_D_, ΔH, ΔS and stoichiometry (n). Binding measurements were performed in triplicate. In a control titration performed with buffer replacing eIF4GII HEAT-1, the heat evolved was constant and less than 0.4 kcal/mol.

### Nuclear Magnetic Resonance (NMR) experiments

NMR chemical shift mapping was used to map interactions made by MNV VPg with the HEAT-1 domain of eIF4GI. Experiments were performed using a Bruker Avance II 800 mHz spectrometer at 298 K. Titration of MNV VPg 104-124 peptide into a solution containing ^15^N labelled untagged eIF4GI HEAT-1 truncated (748-993) was monitored by recording ^1^H-^15^N transverse-relaxation optimized spectroscopy (TROSY) NMR spectra. MNV VPg(104-124) peptide (ChinaPeptides Co., Ltd. Shanghai, China) was dissolved in 50 mM MES pH 6.5, 150 mM NaCl, 5 mM 2-mercaptoethanol and titrated to pH 6.5 using a very small volume of 10 M NaOH. The reference ^1^H-^15^N TROSY spectrum was recorded from a 450 µL sample of 15N eIF4GI HEAT-1 protein at 212 µM dissolved in the same buffer containing 5% D_2_O used as a lock signal. The effects of peptide titration were measured at peptide:protein molar ratios of 0.117, 0.352, 0.587, 0.936, and 1.404 by sequential addition of 1.83 µL of peptide stock solution (6.1 mM). The number of scans recorded in each ^1^H-^15^N TROSY NMR spectra were as follows, reference (16), 0.117 (24), 0.352 (28), 0.587 (56), 0.936 (56), 1.404 (180). The data were transformed using NMRpipe and analysed using NMRview (One Moon Scientific) [62]. The deposited ^1^H-^15^N assignments for eIF4GI HEAT-1 truncated 748-993 (BMRB Entry 18738) was used to analyse the titration [43]. For peak volume analysis the peak list was adjusted to account for slight spectral changes. In addition, clearly overlapped peaks were deleted from the list. The eVolume function of NMRview was used to determine non-overlapped peak volumes.

### Size exclusion chromatography with multi-angle light scattering (SEC-MALLS)

To test for the formation of a ternary complex, untagged eIF4A 1-406, untagged MNV VPg 1-124 and N-terminally His-tagged eIF4GI HEAT-1 truncated (748-993) were mixed together in a total volume of 210 µL with final concentrations of 117 µM, 110 µM, and 132 µM of eIF4A, eIF4G and MNV VPg respectively. The mix was rotated slowly at room temperature for 30 minutes and then concentrated in an Amicon Ultra 500 10,000 MWCO concentrator to 130 µL. This sample was injected onto a S200 10/300 GL column at 0.5 mL/min using a 1260 Infinity HPLC system (Agilent Technologies). Light scattering and refractive index were measured using in-line miniDAWN TREOS and Optilab T-rEX detectors (Wyatt Technology).

Binary complexes (eIF4A:eIF4GI HEAT-1 and MNV VPg: 4GI HEAT-1) and individual protein samples were prepared the same way with the exception that matched buffer replaced one or more of the protein components in the mix. SEC MALS data were analysed by Astra 6 software (Wyatt Technology) using a dn/dc value of 0.185 and weight-averaged molar masses are reported. Peak protein fractions were also analysed by SDS PAGE and Coomassie staining.

### *In vitro* translation assays

MNV1 VPg-linked RNA was isolated from infected RAW 264.7 cells as described previously [27]. For these assays we used capped bicistronic constructs that contain an upstream chloramphenicol acetyl transferase (CAT) open reading frame (ORF) separated from a downstream luciferase (LUC) ORF by the IRES from either porcine teschovirus (PTV) or FMDV, which have been described previously [27]. *In vitro* translation was performed using the Flexi rabbit reticulocyte lysate system (Promega) in the presence of ^35^S-labelled methionine (EasyTag, Perkin Elmer). Translation reactions were performed essentially according to the manufacturer’s protocol but scaled down to 12.5 µL and with the inclusion of 100mM KCl. They were pre-incubated for 15 minutes with 3.5-28 µM purified protein GST-MNV VPg(102-124) (wild-type or F123A mutant) and then initiated by addition of either 25 ng/µL *in vitro* transcribed bicistronic RNA or 16 ng/µL MNV1 VPg-linked RNA to give a final reaction volumes of 12.5 µL. The reactions were incubated at 30°C for 90 min and terminated by addition of an equal volume of Tris buffer containing 10 mM ethylenediaminetetraacetic acid (EDTA) and 100 ng/mL RNase A. Translated proteins were re-suspended in SDS PAGE sample buffer and resolved on a 12.5 % polyacrylamide gel. Protein synthesis levels were quantified by autoradiography. The intensity of each band was quantified with reference to the value obtained in the absence of recombinant protein.

### Lentivirus vector particle production and transduction

Lentivirus vectors were used to test intracellular effects of MNV VPg peptide sequences expressed as C-terminal fusions with the fluorescent mCherry protein tag (see above). Vesicular stomatitis virus G-protein-pseudotyped lentiviral particles were generated by transient transfection of 293T cells grown in 6-wells plates using 1.25 µg lentiviral vector, 0.63 µg pMDLg/pRRE (Addgene #12251), 0.31 µg pRSV-Rev (Addgene#12253) and 0.38 µg pMD2.G (Addgene#12259) per well using published protocols [63]. Parental BV2 cells were transduced with lentiviral supernatants and incubated for 48 h. Transduced cells were then selected on the basis of their resistance to Hygromycin B at a concentration of 200 µg/mL and sub-cultured for three additional passages. Cell populations expressing equal level of mCherry fluorescence were then enriched by FACS prior to infection with MNV1 or immunoprecipitation of mCherry-VPg protein (see below).

### Co-immunoprecipitation experiments with mCherry-VPg

Approximately 107 BV2 transduced cells were washed with cold phosphate-buffered saline (PBS) and harvested in 200 µL lysis buffer (Tris-HCl 10 mM pH 7.5, NaCl 150 mM, NP40 0.5 %, EDTA 0.5 mM, PMSF 1 mM and 1x protease inhibitors (Calbiochem)). Cells were disrupted by up and down pipetting and lysates were centrifuged at 12,000 g for 10 min at 4°C. Clarified supernatants were diluted to 500 µL with lysis buffer. RFP-Trap^®^_A agarose beads (Chromotek GmbH, Cat No: RTA-20) were added and the lysates were incubated for 2 h at 4°C under gentle agitation. The beads were washed 5 times with lysis buffer and proteins were eluted with Laemmli sample buffer. Immunoprecipitated proteins were detected using SDS PAGE and immunoblot analysis. As a control of protein expression in cells used for co-immunoprecipitation experiments, total cell lysate corresponding to 5 % of the input used for immunoprecipitation was also analysed by immunoblot.

Antibodies used were obtained from a variety of sources. Mouse monoclonal anti-eIF4E (A-10) (Cat No: sc-271480), rabbit polyclonal anti-eIF4G (H-300) (4GI) (cat No: sc-11373) and goat polyclonal anti-eIF4AI (N-19) (Cat No: sc-14211) were obtained from Santa Cruz Biotechnology Inc. Rabbit polyclonal anti-eIF3D was obtained from ProteinTech (Cat No: 10219-1-AP). Mouse monoclonal anti-GAPDH was obtained from Ambion (Cat No: AM4300). Rat monoclonal anti-RFP (clone 5F8) was obtained from Chromotek GmbH. Rabbit polyclonal anti-PABP1 was obtained from Cell Signaling Technology (Cat No: #4992).

### Measurement of viral replication

The effect of excess added MNV VPg peptide sequences on MNV replication was tested using cell-penetrating peptides and ectopic expression of mCherry VPg.

We used synthetic FITC-labelled peptides containing cell-penetrating sequences from Bac7 or human Immunodeficiency Virus (HIV) TAT fused to residues 102-124 of MNV VPg (sourced from ChinaPeptides Co. Ltd., Shanghai, China). F123A mutant variants of both peptides (which bind much less well to eIF4G) were also used. The peptide sequences were as follows: Bac-WT: FITC-PRPLPFPRPGNVVGPSWADDDRQVDYGEKINFE; Bac-F123A: FITC-PRPLPFPRPGNVVGPSWADDDRQVDYGEKINAE; TAT-WT: FITC-GRKKRRQRRRPQNVVGPSWADDDRQVDYGEKINFE; TAT-F123A: FITC-GRKKRRQRRRPQNVVGPSWADDDRQVDYGEKINAE. The identity and purity (>95%) of the labelled peptides were assessed by mass spectrometry and reverse-phase liquid chromatography. All peptides were dissolved in dimethyl sulfoxide (DMSO) at a concentration of 10 mM and stored at 80°C.

Naïve BV2 cells were treated with peptides at a concentration of 100 µM in culture complete medium for 150 min before being infected with MNV1 at low MOI (0.01 TCID_50_ units/cell) in the presence of peptides. Untreated cells were cultured in the same conditions using DMSO. After the indicated period of time, total cell RNA was extracted using a GenElute Mammalian Total RNA Miniprep kit (Sigma) and normalised to a standard concentration before being reverse transcribed using random hexamers and MuMLV RT enzyme (Promega). SYBR green-based quantitative PCR was performed using MNV1-specific primers (Supplementary Table S2). Each sample was measured in biological triplicate and compared to a standard curve. Additional non-template and non-reverse transcriptase samples were analysed as negative controls. Data were collected using a ViiA 7 Real-Time PCR System (Applied Biosystems).

Measurements of viral replication in BV2 cells ectopically expressing mCherry-VPg fusion proteins or mCherry alone (prepared by lentiviral transduction as described above) were performed using a similar experimental setup.

## Acknowledgements

We thank Xulin Liu and Stephen Hare for technical assistance, and Trevor Sweeney for critical reading of the manuscript.

---

**Figure S1.**
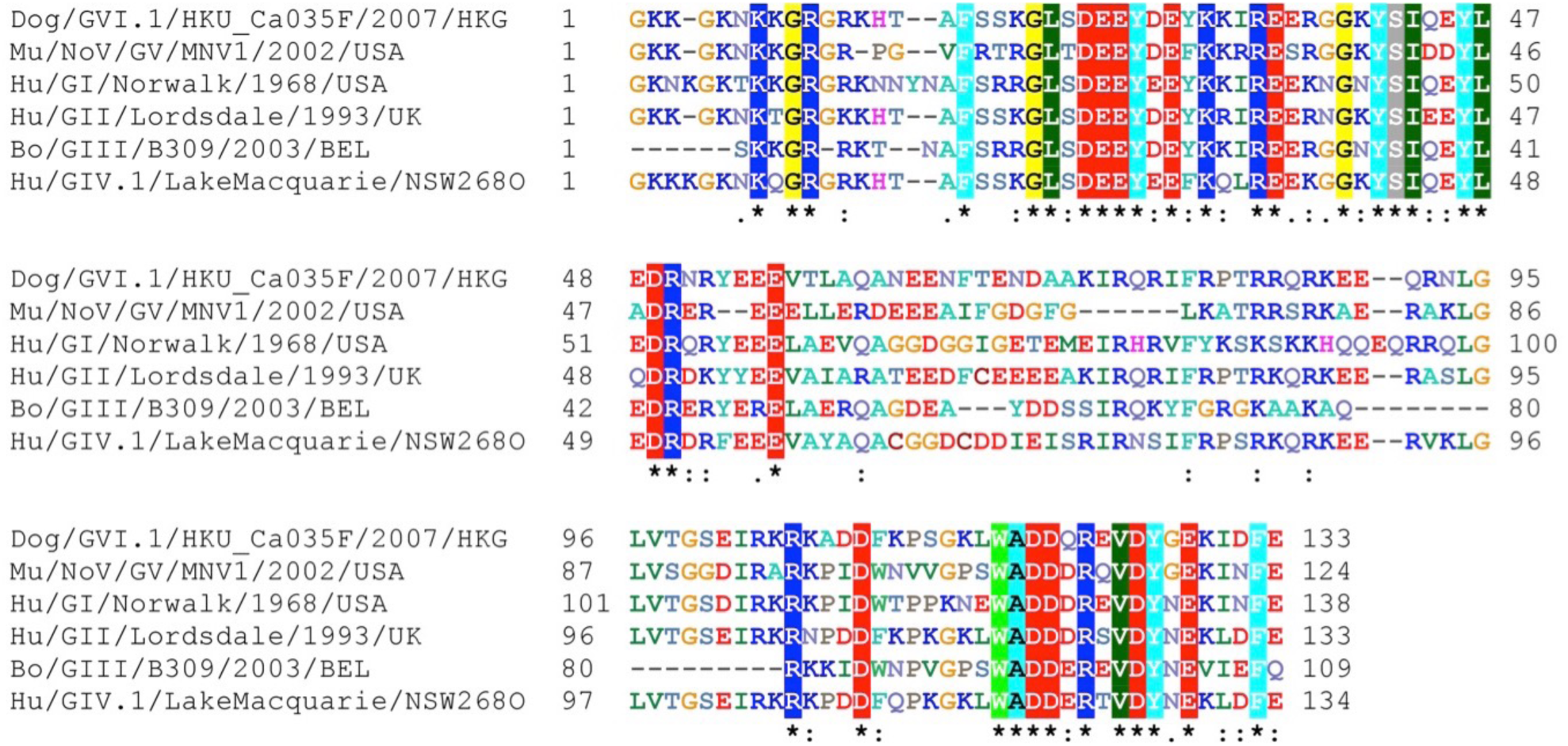
**Supplementary Figure 1: Amino acid sequence alignments of representative sequences for VPg of all 6 genogroups of Norovirus (GI-GIV).** The representative strains used in the alignments are GI Hu/GI/Norwalk/1968/US, (NCBI accession AAC64602), GII Lordsdale virus Hu/GII/Lordsdale/1993/UK (NCBI accession P54634), GIII Bo/GIII/B309/2003/BEL (NCBI accession ACJ04905.1),Hu/GIV.1/LakeMacquarie/NSW268O (NCBI accession number AFJ21375), Mu/NoV/GV/MNV1/2002/ USA (NCBI accession ABU55564.1), GVI dog/GVI.1/HKU_Ca035F/2007/HKG (NCBI accession FJ692501). Sequence alignment was performed by ClustalW [64] and BioEdit (http://www.mbio.ncsu.edu/BioEdit/bioedit.html).

**Figure S2.**
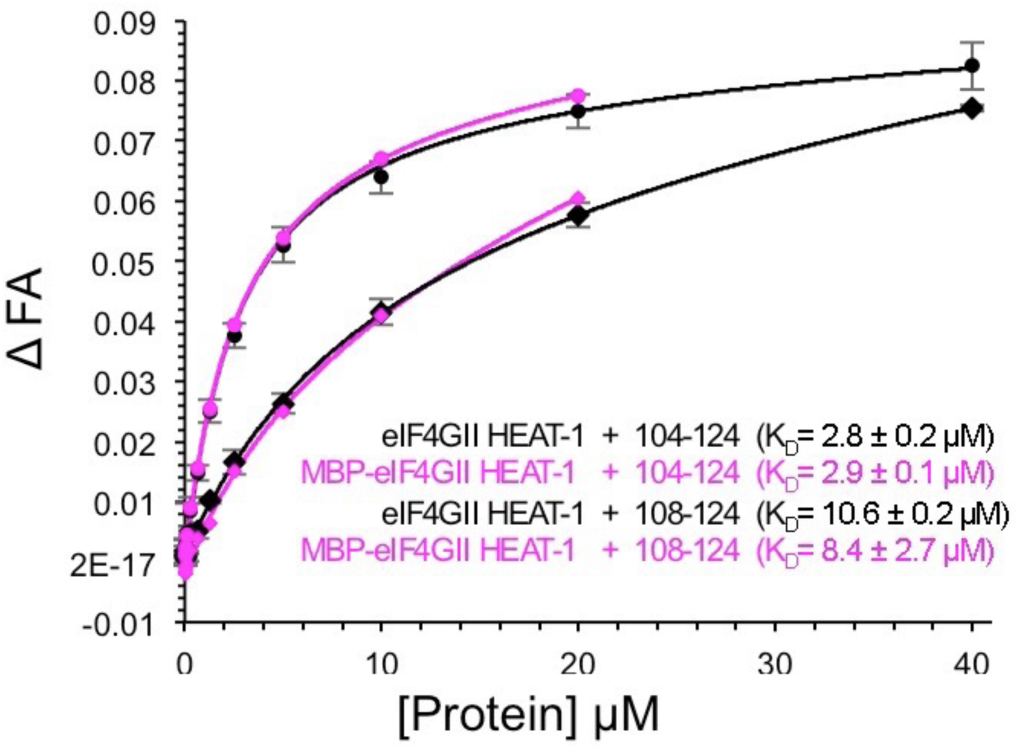
**Supplementary Figure 2: Comparison of binding of eIF4GII HEAT-1 and MBP-eIF4GII HEAT-1 to MNV VPg C-terminal peptides.** FITC-labelled MNV VPg(104-124) and MNV VPg(108-124) peptides were used in fluorescence anisotropy binding assays with the HEAT-1 domain of eIF4GII (751-1009) in order to determine the K_D_ of the interaction. ΔFP, the normalised change in fluoresce anisotropy (relative to a no protein control) is plotted against protein concentration. Where appropriate (N>1), error bars indicate the standard deviation in the mean ΔFP value observed. The assays were performed with untagged eIF4GII HEAT-1 (black) and an MBP-tagged version (purple). The data generated using the untagged protein were fit using GraphPad Prism to a single-site binding model. Solid line: MNV VPg(104-124); dashed line - MNV VPg(108-124).

**Figure S3.**
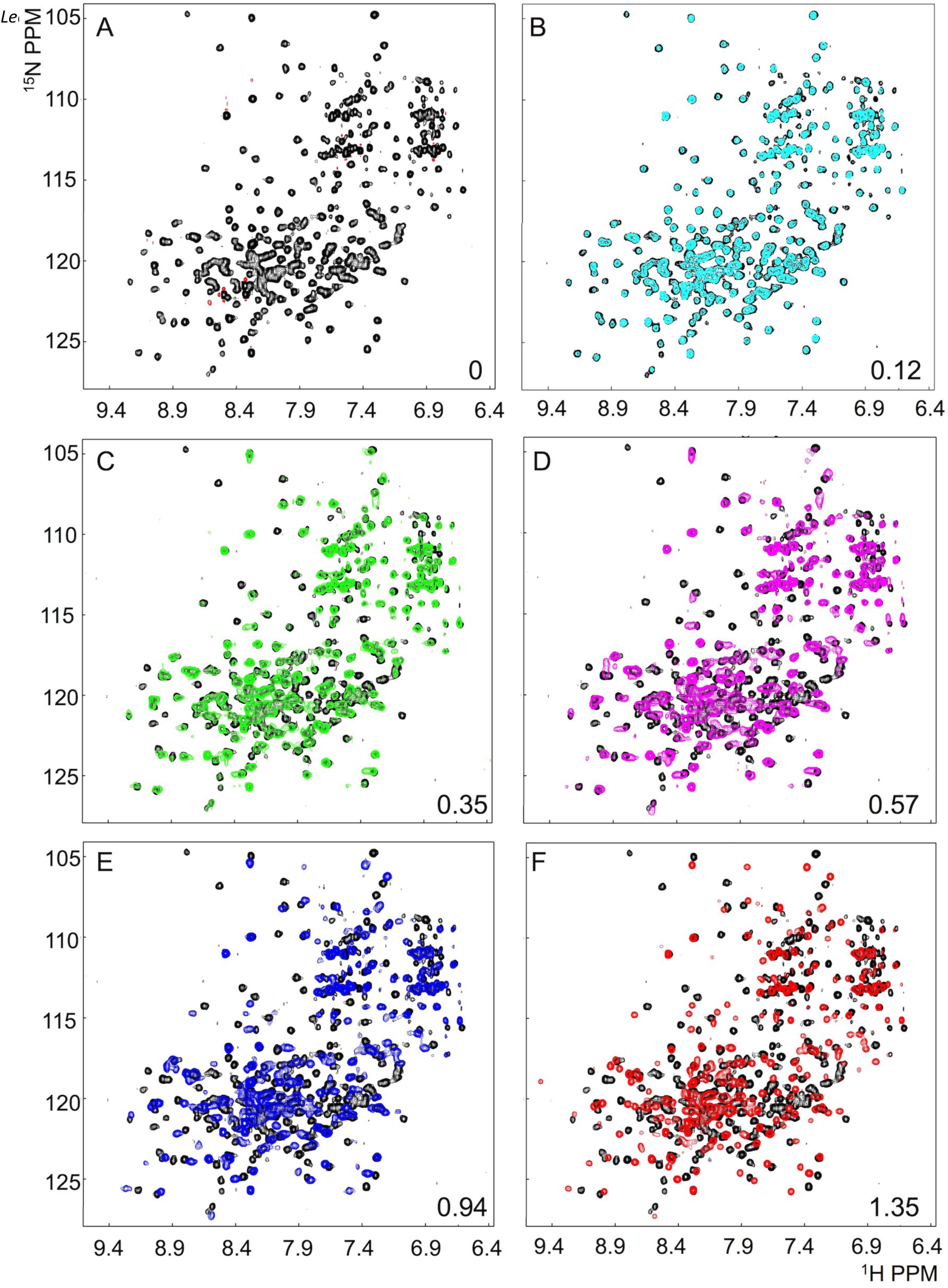
**Supplementary Figure 3: NMR analysis of binding of MNV VPg(104-124) to eIF4GI.** ^1^H-^15^N TROSY HSQC spectra obtained on titration of unlabelled MNV VPg(104-124) into 212 µM 15N-labelled eIF4GI HEAT-1 (748-993). (A) Reference spectrum obtained in the absence of MNV VPg(104-124). (B-F) Spectra obtained in the presence of (B) 0.12, (C) 0.35, (D) 0.57, (D) 0.94 and (F) 1.35 molar equivalents of MNV VPg(104-124) peptide superposed on the reference spectrum. The molar equivalents of MNV VPg(104-124) peptide and the number of scans used to obtain the spectrum (which was increased as the average signal intensity decreased) each point in the titration.

**Figure S4.**
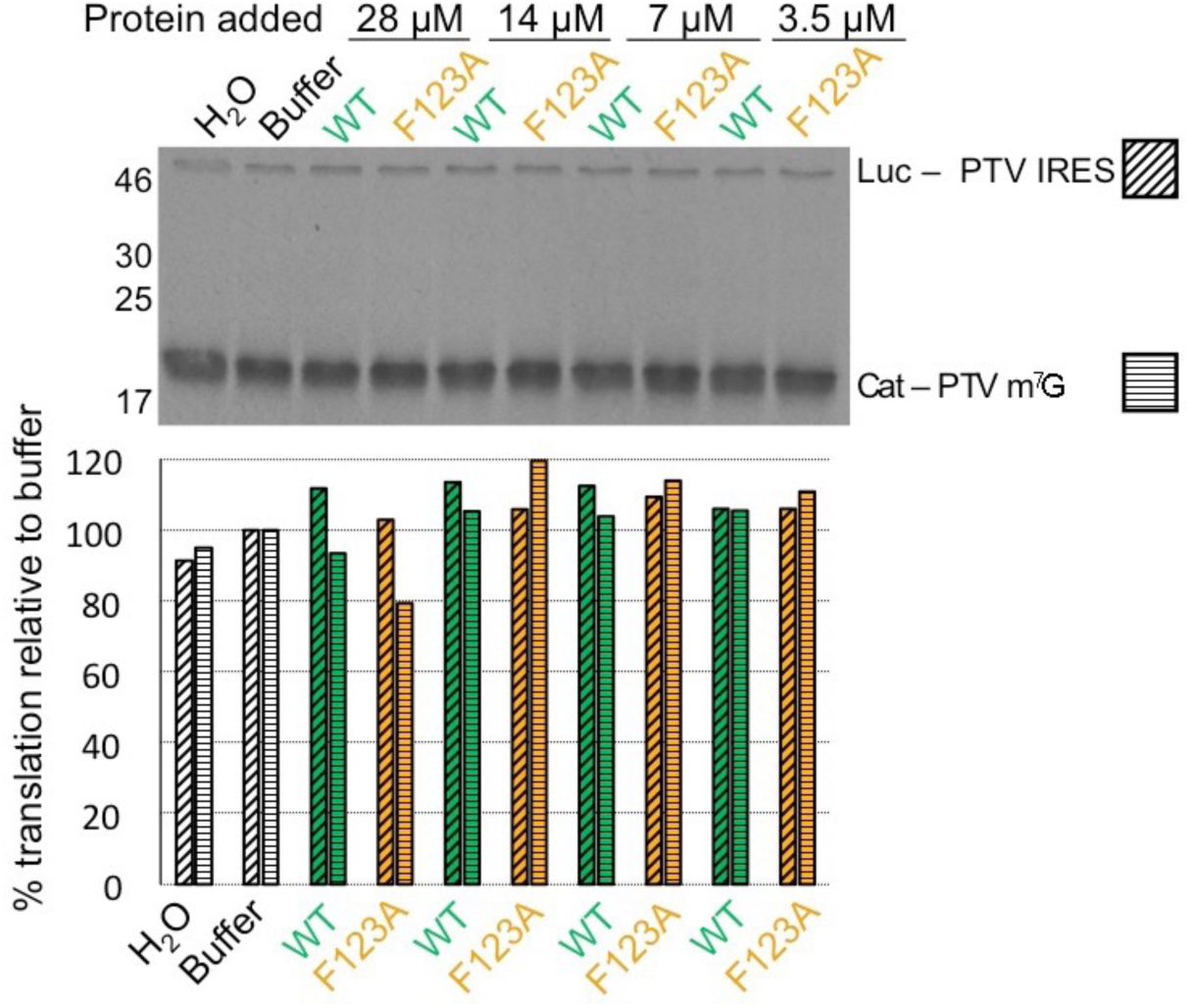
**Supplementary Figure 4: GST-MNV VPg(102-124) does not inhibit cap-dependent or PTV IRES-dependent translation.** *In vitro* translation reactions were performed in the presence of increasing concentrations of GST-MNV VPg(102-124) WT protein or the GST-MNV VPg(102-124) F123A mutant that binds much less well eIF4G. Protein synthesis was monitored by autoradiography of SDS PAGE analysis of incorporation of ^35^S-methionine in translation reactions. Top panel: Effect of exogenous GST-MNV VPg(102-124) proteins on translation from capped bi-cistronic mRNA constructs containing the PTV IRES between the first (CAT) and second (Luc) cistrons; bottom panel: quantitative analysis of the level of ^35^S-methionine incorporation.

**Figure S5.**
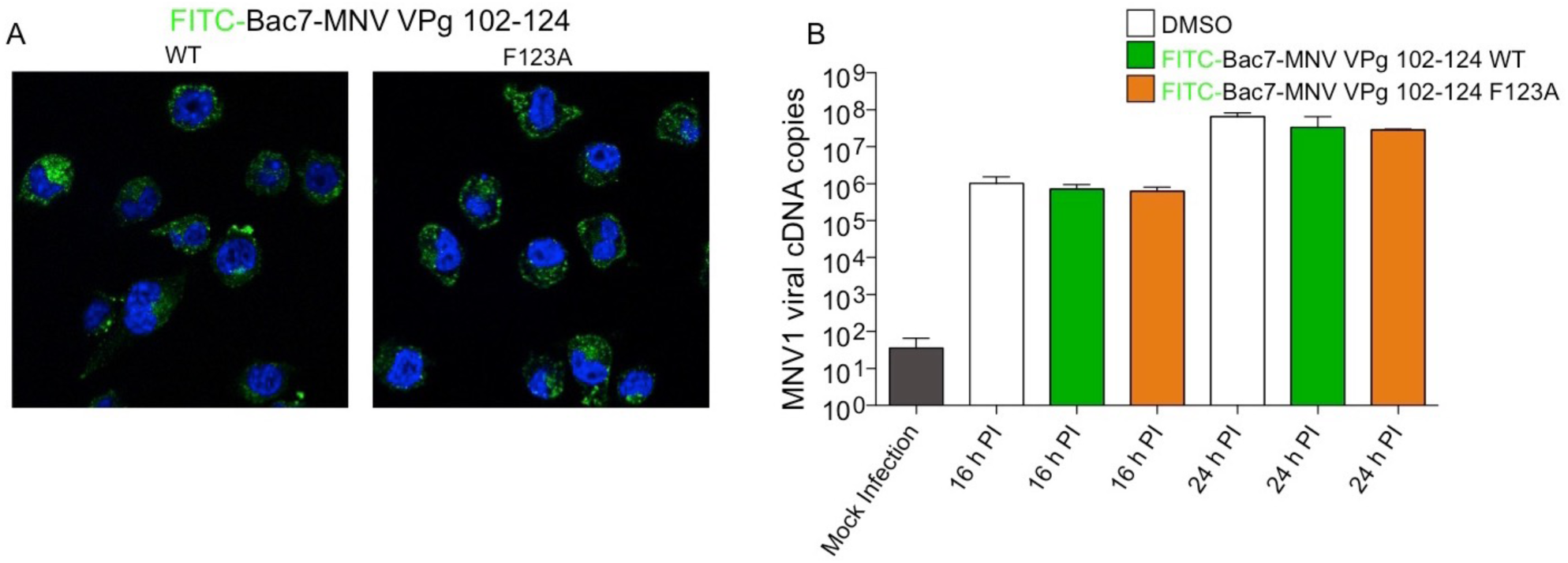
**Supplementary Figure 5: Cell penetrating FITC-Bac7-MNV VPg(102-124) peptides do not inhibit MNV infection of BV2 cells.** (A) Fluorescence microscopy analysis of cell penetration of the FITC-Bac7-MNV VPg 102-124 peptides. The images shown are merged images of DAPI stained nuclear DNA (blue) and wild-type or F123A versions of the cell penetrating peptides (green). (B) Time course of BV2 infection (MOI 0.01 TCID_50_ units/cell) with MNV1 following pre-treatment for 150 minutes with 100 µM of wild-type or F123A versions of the cell penetrating peptides prior to infection. The progress of infection was monitored by RT-PCR analysis (as in Fig 8).

## Supplementary Materials and Methods – Specific adjustments to purification protocols

### MNV VPg 1-124 WT and mutants used in mutational mapping of the eIF4G-VPg interaction

Six of the hexa-His-tagged MNV VPg 1-124 protein constructs (wild-type, F123A, ^120^KIN→^120^ESA, ^116^DYGE→^116^RAPK, ^112^DRQV→^112^REAS, and ^108^WADD→^108^APRR) used to delineate the extent of the eIF4GI binding site in cobalt affinity pull-down assays (presented in Fig 2B) were initially purified on TALON resin essentially as described in Materials and Methods. However, following overnight dialysis in 50 mM HEPES pH 7.3, 300 mM NaCl, 2 mM 2-mercaptoethanol, two of the mutant proteins – 116DYGE→116RAPK and 108WADD→108APRR – were found to have OD260/280 ratios of 1.51 and 1.95 respectively. These were unusually high and suggested significant nucleic acid contamination. In order to remove any bound nucleic acids the proteins were diluted approximately ten-fold with phosphate buffer (15.8 mM Na_2_HPO_4_, 34.2 mM NaH_2_PO_4_, pH 6.5, 300 mM NaCl, 1 mM DTT) and protamine sulfate was added to a concentration of 1 mg/mL. Precipitated nucleic acids were removed by centrifugation at 29,000g for 20 minutes. The protein in the supernatant was then rebound to TALON cobalt resin (Clontech) for 1 h with slow rotation, after which the slurry was applied to a gravity flow column and the resin washed with 75 mL of phosphate buffer. The protein was eluted with 10 mL of phosphate buffer supplemented with 100 mM imidazole. Eluted fractions were dialysed for 13 hours against 4 L phosphate buffer and the proteins concentrated and frozen as described in the Material and Methods section.

The MNV VPg ^104^VGPS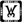^104^KKAH mutant and 103GSGSGS purifications were modified based on the purifications of the other mutants. These were purified as per the Materials and Methods. 50 mM sodium phosphate pH 6.5 (15.8 mM Na_2_HPO_4_, 34.2 mM NaH_2_PO_4_), 300 mM NaCl, 1 mM DTT was used as the buffer for both purification buffer and dialysis. The final A260/280 of the WT and all the mutant MNV VPg 1-124 proteins was 0.61 or less.

### eIF4GI HEAT-1 truncated (748-993) wild-type and mutants used in mutational mapping of the eIF4G-VPg interaction

The wild-type and mutant eIF4GI HEAT-1 (748-993) proteins were initially purified, to completion, as described in Materials and Methods. However the high OD260/280 ratio of some of the mutant proteins was suggestive of nucleic acid contamination (D919R – 1.6, L939A – 1.04, H918A – 1.145, K901M-E914R – 1.15, L897A – 0.84). Therefore the purified proteins were thawed and incubated with 1 µM MgCl_2_ and 387 U benzonase nuclease (Sigma) for 70-90 minutes (typically 1 U benzonase per 0.02 mg protein). The mixture was then subjected to size-exclusion chromatography on a Superdex 75 10/300 GL column equilibrated with 10.1 mM Na_2_HPO_4_, 1.8 mM KH_2_PO_4_, pH 7.2, 136.9 mM NaCl, 2.7 mM KCl, and 1 mM tris(2-carboxyethyl)phosphine hydrochloride (TCEP.HCl). Peak fractions were pooled, concentrated and stored frozen. The final OD260/280 ratios were all under 0.7.

**Supplementary Table 1.**
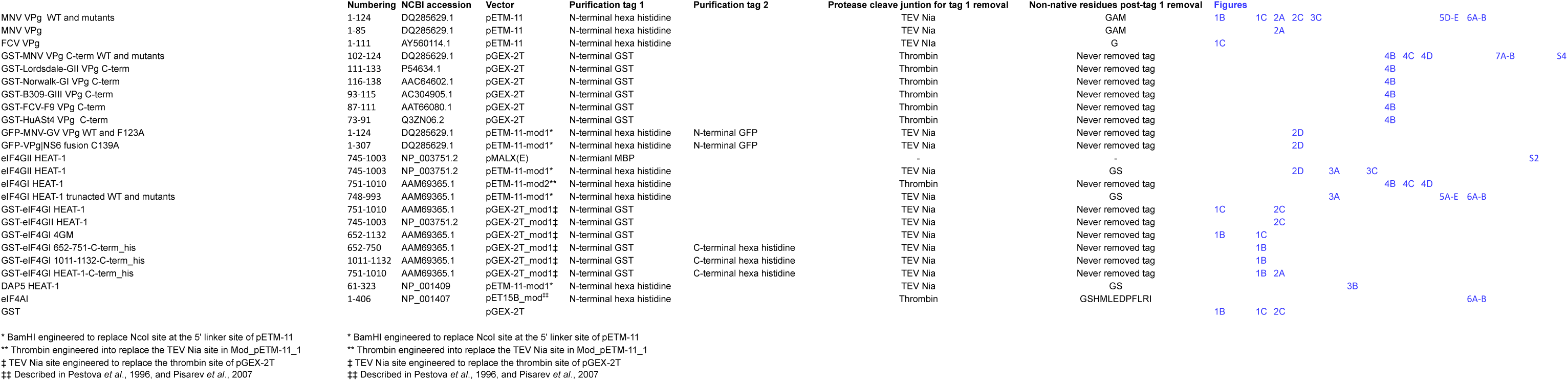
Construct,Details

**Supplementary Table 2.**
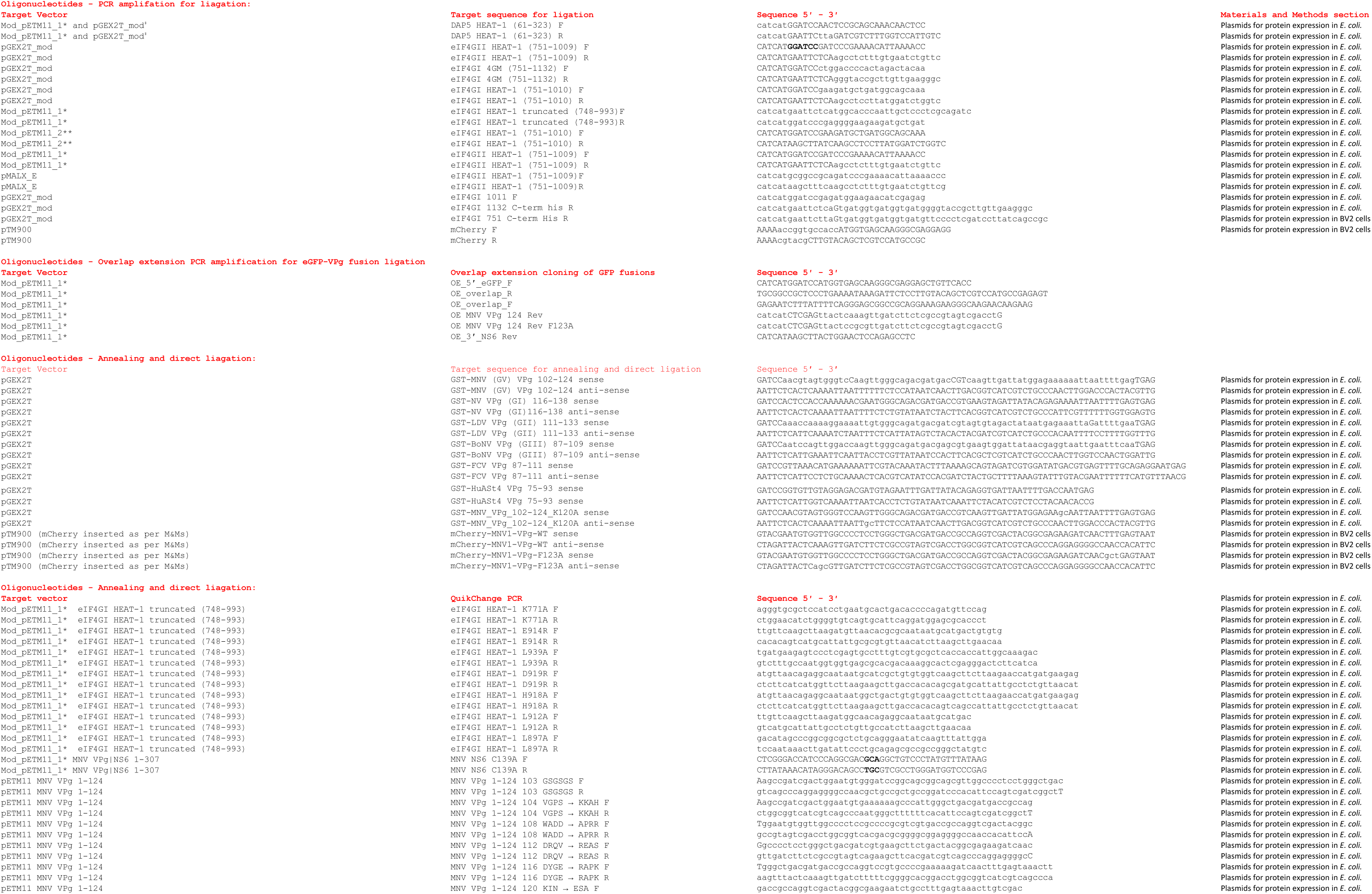
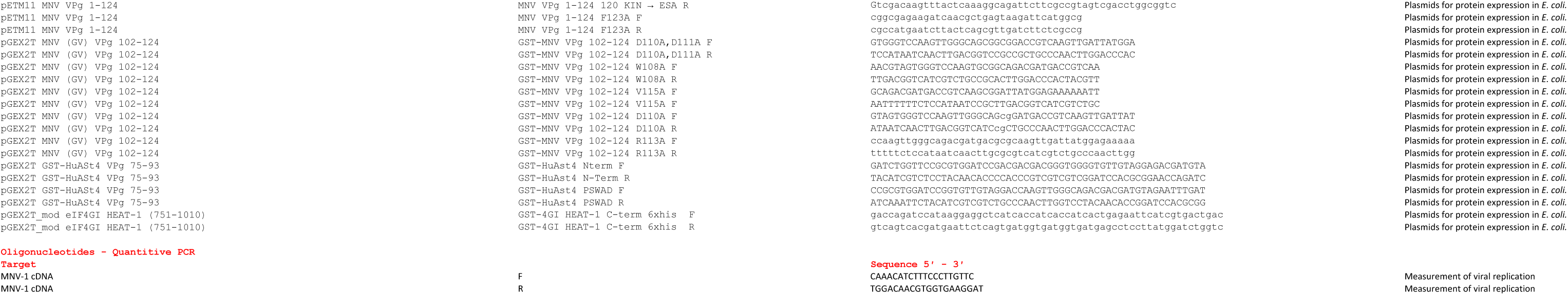
Primers

**Supplementary Table 3.** Expression Details

|  | Numbering | NCBI accession | Vector | *E. coli* cell type | [IPTG] mM | Expression temp (° C) | RPM | Expression time (hours) | Other details |
| --- | --- | --- | --- | --- | --- | --- | --- | --- | --- |
| MNV VPg WT and mutants | 1-124 | DQ285629.1 | pETM-11 | BL21-Rosetta(DE3) | 1 | 20 | 220 | 4 | Chilled on ice for 5 min before induction |
| FCV VPg | 1-111 | AY560114.1 | pETM-11 | BL21-Rosetta(DE3) | 1 | 20 | 220 | 4 | Chilled on ice for 5 min before induction |
| GST-MNV VPg C-term WT and mutants | 102-124 | DQ285629.1 | pGEX-2T | BL21-CodonPlus(DE3) RIPL | 1 | 37 | 220 | 4 |  |
| GST-LDV VPg C-term | 111-133 | P54634.1 | pGEX-2T | BL21-CodonPlus(DE3) RIPL | 1 | 37 | 220 | 4 |  |
| GST-NV VPg C-term | 116-138 | AAC64602.1 | pGEX-2T | BL21-CodonPlus(DE3) RIPL | 1 | 37 | 220 | 4 |  |
| GST-BoNV VPg C-term | 93-115 | AC304905.1 | pGEX-2T | BL21-CodonPlus(DE3) RIPL | 1 | 37 | 220 | 4 |  |
| GST-FCV VPg C-term | 87-111 | AAT66080.1 | pGEX-2T | BL21-CodonPlus(DE3) RIPL | 1 | 37 | 220 | 4 |  |
| GST-HuASt4 VPg C-term WT and mutants | 73-91 | Q3ZN06.2 | pGEX-2T | BL21-CodonPlus(DE3) RIPL | 1 | 37 | 220 | 4 |  |
| GFP-MNV VPg WT and F123A | 1-124 | DQ285629.1 | pETM-11-mod1* | BL21-CodonPlus(DE3) RIPL | 1 | 20 | 220 | 4 | Chilled on ice for 5 min before induction |
| GFP-VPg\|NS6 fusion C139A | 1-307 | DQ285629.1 | pETM-11-mod1* | BL21-CodonPlus(DE3) RIPL | 1 | 20 | 220 | 4 | Chilled on ice for 5 min before induction |
| eIF4GII HEAT-1 | 745-1003 | NP_003751.2 | pMALX(E) | BL21-CodonPlus(DE3) RIPL | 1 | 37 | 220 | 3 |  |
| eIF4GII HEAT-1 | 745-1003 | NP_003751.2 | pETM-11-mod1* | BL21-CodonPlus(DE3) RIPL | 1 | 30 | 220 | 4 |  |
| eIF4GI HEAT-1 | 751-1010 | AAM69365.1 | pETM-11-mod2** | BL21-CodonPlus(DE3) RIPL | 1 | 20 | 220 | 7 |  |
| eIF4GI HEAT-1 trunacted WT and mutants | 748-993 | AAM69365.1 | pETM-11-mod1* | BL21-CodonPlus(DE3) RIPL | 1 | 30 | 220 | 4 |  |
| GST-eIF4GI HEAT-1 | 751-1010 | AAM69365.1 | pGEX-2T_mod1‡ | BL21-Rosetta(DE3) | 1 | 37 | 220 | 4 |  |
| GST-eIF4GII HEAT-1 | 745-1003 | NP_003751.2 | pGEX-2T_mod1‡ | BL21-Rosetta(DE3) | 1 | 37 | 220 | 4 |  |
| GST-eIF4GI 4GM | 652-1132 | AAM69365.1 | pGEX-2T_mod1‡ | BL21-CodonPlus(DE3) RIPL | 1 | 30 | 220 | 4 |  |
| GST-eIF4GI 652-751-C-term_his | 652-750 | AAM69365.1 | pGEX-2T_mod1‡ | BL21-CodonPlus(DE3) RIPL | 1 | 30 | 220 | 4 |  |
| GST-eIF4GI 1011-1132-C-term_his | 1011-1132 | AAM69365.1 | pGEX-2T_mod1‡ | BL21-CodonPlus(DE3) RIPL | 1 | 30 | 220 | 4 | Chilled on ice for 5 min before induction |
| GST-eIF4GI HEAT-1-C-term_his | 751-1010 | AAM69365.1 | pGEX-2T_mod1‡ | BL21-CodonPlus(DE3) RIPL | 1 | 30 | 220 | 4 |  |
| DAP5 HEAT-1 | 61-323 | NP_001036024.3 | pETM-11-mod1* | BL21-CodonPlus(DE3) RIPL | 1 | 30 | 220 | 4 |  |
| GST-DAP5 HEAT-1 | 61-323 | NP_001036024.3 | pGEX-2T_mod1‡ | BL21-CodonPlus(DE3) RIPL | 1 | 30 | 220 | 4 |  |
| eIF4A | 1-406 | NP_001407 | pET15B_mod‡‡ | BL21-CodonPlus(DE3) RIPL | 1 | 30 | 220 | 4.5 |  |
| GST |  |  | pGEX-2T | BL21-Rosetta(DE3) | 1 | 37 | 220 | 4 |  |

\* BamHI engineered to replace NcoI site at the 5' linker site of pETM-11 \*\* Thrombin engineered into replace the TEV Nia site in Mod\_pETM-11\_1 ‡ TEV Nia site engineered to replace the thrombin site of pGEX-2T ‡‡ Described in Pestova *et al.*, 1996, and Pisarev *et al.*, 2007.

**Supplementary Table 4.**
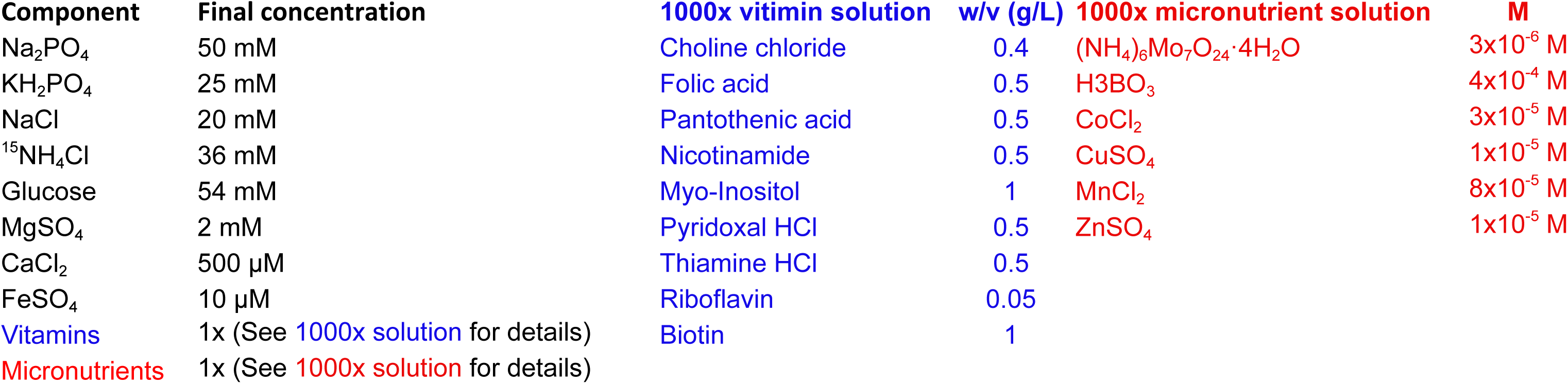
NMR Solutions

| Component | Final concentration | 1000x vitamin solution | w/v (g/L) | 1000x micronutrient solution | M |
| --- | --- | --- | --- | --- | --- |
| Na<sub>2</sub>PO<sub>4</sub> | 50 mM | Choline chloride | 0.4 | (NH<sub>4</sub>)<sub>6</sub>Mo<sub>7</sub>O<sub>24</sub>·4H<sub>2</sub>O | 3x10<sup>-6</sup> M |
| KH<sub>2</sub>PO<sub>4</sub> | 25 mM | Folic acid | 0.5 | H<sub>3</sub>BO<sub>3</sub> | 4x10<sup>-4</sup> M |
| NaCl | 20 mM | Pantothenic acid | 0.5 | CoCl<sub>2</sub> | 3x10<sup>-5</sup> M |
| <sup>15</sup>NH<sub>4</sub>Cl | 36 mM | Nicotinamide | 0.5 | CuSO<sub>4</sub> | 1x10<sup>-5</sup> M |
| Glucose | 54 mM | Myo-Inositol | 1 | MnCl<sub>2</sub> | 8x10<sup>-5</sup> M |
| MgSO<sub>4</sub> | 2 mM | Pyridoxal HCl | 0.5 | ZnSO<sub>4</sub> | 1x10<sup>-5</sup> M |
| CaCl<sub>2</sub> | 500 µM | Thiamine HCl | 0.5 |  |  |
| FeSO<sub>4</sub> | 10 µM | Riboflavin | 0.05 |  |  |
| Vitamins | 1x (See 1000x solution for details) | Biotin | 1 |  |  |
| Micronutrients | 1x (See 1000x solution for details) |  |  |  |  |

**Supplementary Table 5.**
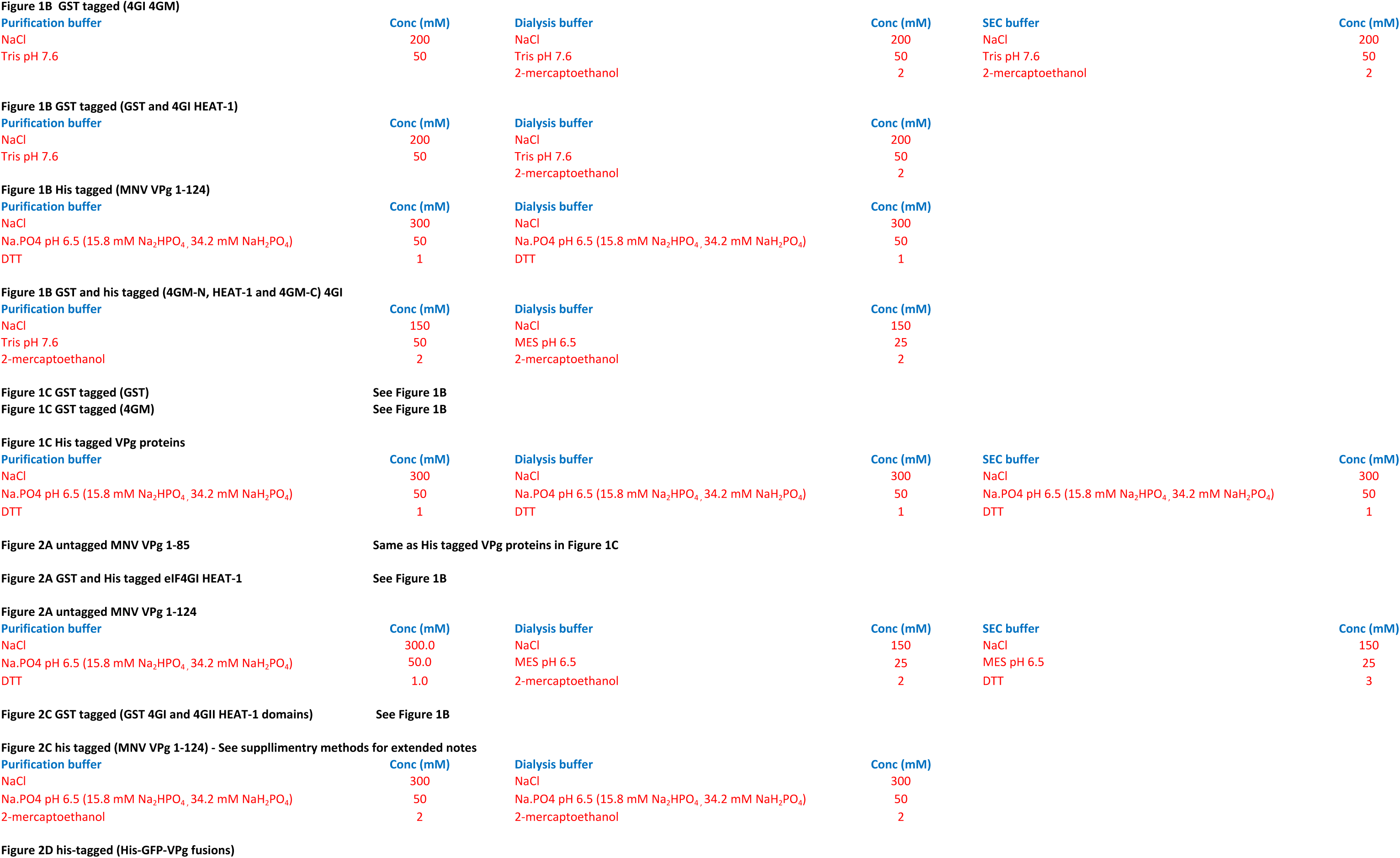
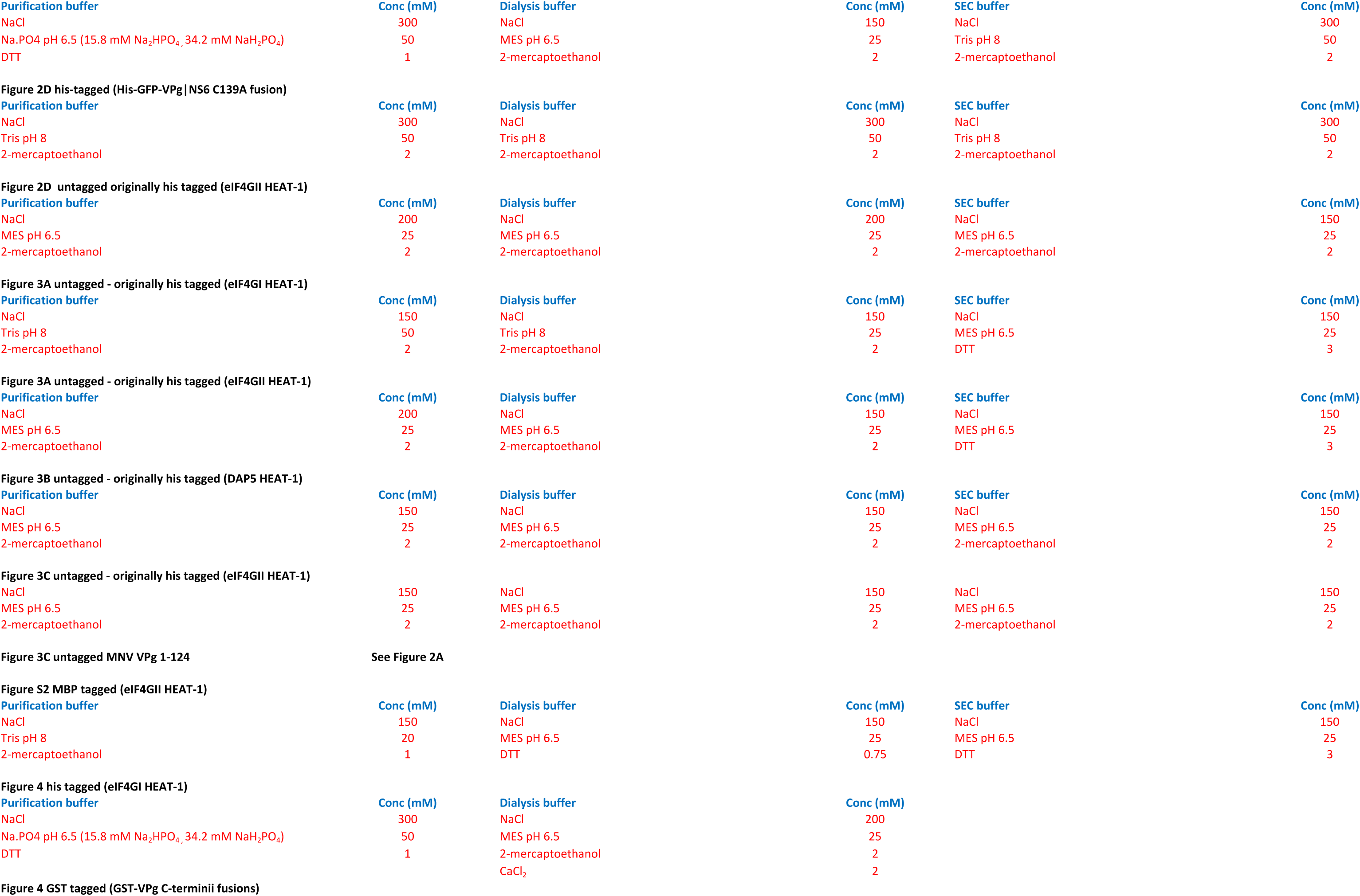
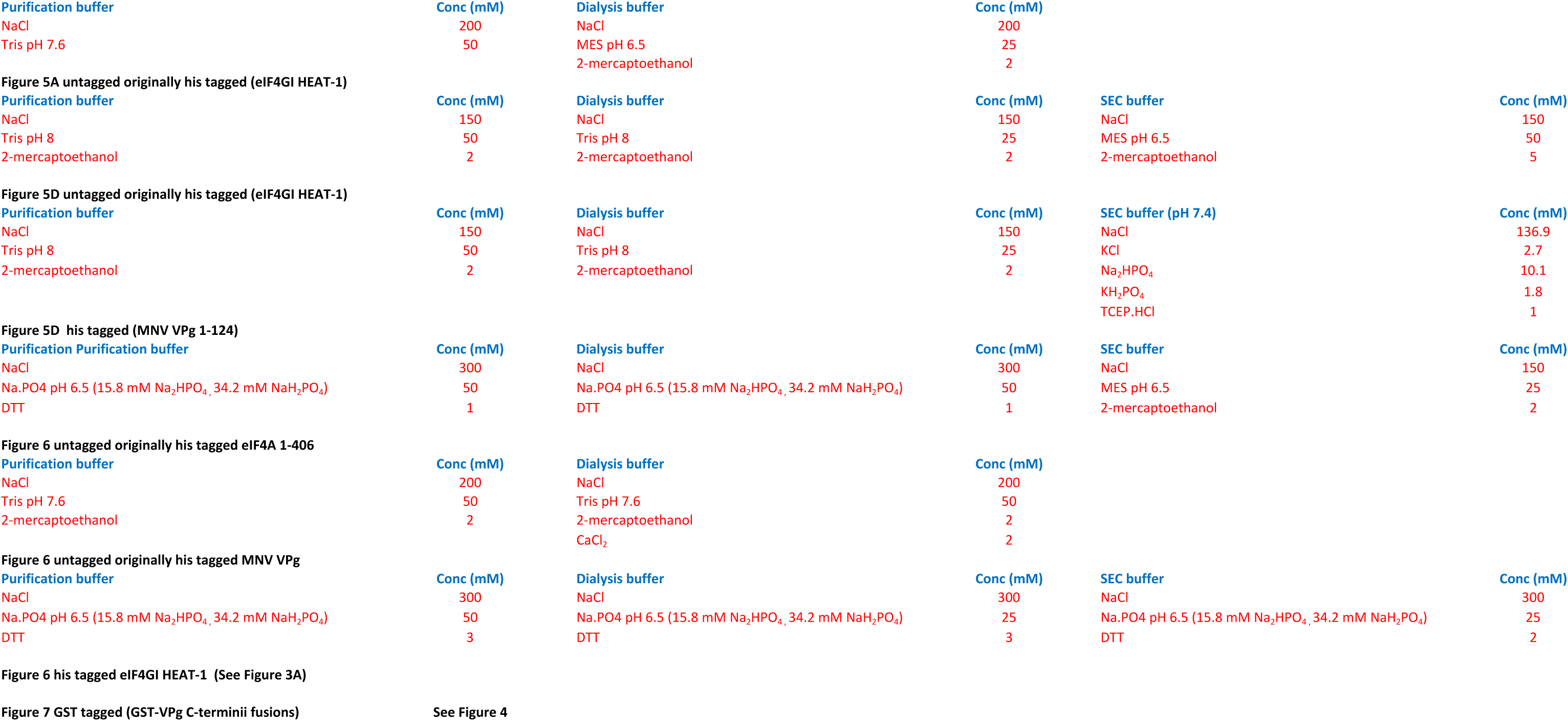
Buffers

